# SpatialESD: spatial ensemble domain detection in spatial transcriptomics

**DOI:** 10.1101/2025.09.11.675735

**Authors:** Hongyan Cao, Gaiqin Liu, Jingyi Xia, Runle Chen, Tong Wang, Xiaoling Yang, Ruiling Fang, Yanhong Luo, Ping Zeng, Hongmei Yu, Yanbo Zhang, Yuehua Cui

## Abstract

Spatial transcriptomics (ST) measures gene expression while preserving spatial context within tissues. One of the key tasks in ST analysis is spatial domain detection, which remains challenging due to the complex structure of ST data and the varying performance of individual clustering methods. To address this, we propose SpatialESD, a Spatial EnSemble Domain detection method that integrates results from different spatial domain detection methods to improve spatial domain detection. SpatialESD captures both direct co-occurrence patterns and multiscale indirect relationships between clusters, improving the robustness and accuracy of spatial domain detection. We evaluated SpatialESD on simulated datasets and multiple 10x Visium spatial transcriptomics datasets, including human brain, breast cancer, and ovarian cancer samples. The results show that SpatialESD consistently outperforms individual methods and the existing EnSDD ensemble method in terms of clustering accuracy and stability. Based on the identified domains, we further detected region-specific differentially expressed genes and performed trajectory and cell-cell interaction analyses. These results reveal spatial patterns of gene expression and cellular communication, offering insights into tissue organization and disease mechanisms. Overall, SpatialESD provides a reliable and effective solution for spatial domain detection in ST data and facilitates downstream biological discovery.

**Graphical Abstract:** 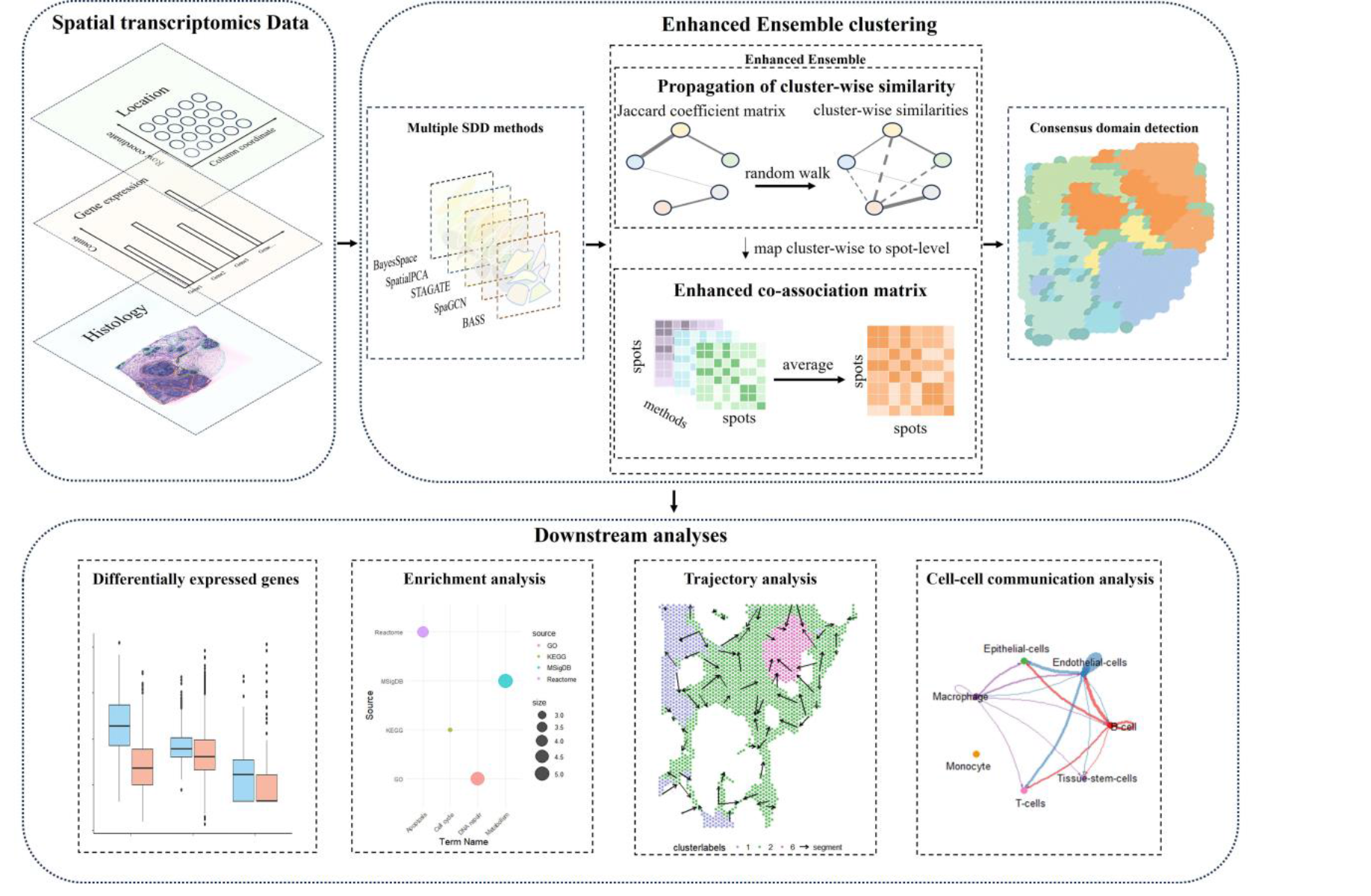

## 1 Introduction

Spatial transcriptomics (ST) is an innovative genomic technology that enables high-throughput transcriptome profiling within tissue sections, while preserving spatial location information ^[1,2]^. These technologies can be broadly classified into imaging-based and sequencing-based methods ^[3]^, each with its own advantages in spatial transcriptomic analysis. Imaging-based methods, such as in situ hybridization (ISH) and in situ sequencing (ISS), rely on fluorescent labeling to detect gene expression patterns, whereas sequencing-based methods, such as Slide-seq and 10x Visium, utilize spatial barcoding technology to perform high-throughput transcriptome profiling at specific regions of tissue sections.

A crucial step in ST studies is the detection of spatial domains, which are regions that exhibit spatial coherence in both gene expression and histological structure ^[4]^. However, the additional dimension of spatial information, which increases data volume and complexity, presents significant challenges for spatial domain detection ^[5]^. Traditional clustering algorithms, such as K-means and Louvain ^[6]^, relying solely on gene expression data, often lead to clusters that do not align well with the spatial organization of tissue sections ^[7]^. Recently, several spatial clustering methods have been developed to enhance spatial domain detection by incorporating spatial coordinates from ST data, thereby accounting for gene expression’s spatial dependencies. These methods can be further divided into two categories ^[8]^: 1) Statistical methods such as BayesSpace ^[9]^, BASS ^[10]^, DR.SC ^[11]^ and SpatialPCA ^[12]^, which model adjacency-based similarity within specialized frameworks to detect clusters. For instance, BayesSpace ^[9]^ utilizes the Potts model as a spatial prior, assuming that adjacent spots are more likely to belong to the same spatial domain; and 2) Graph-based deep learning methods such as STAGATE ^[13]^, SpaGCN ^[4]^ and CCST ^[14]^, primarily utilize graph neural networks to extract latent spot features before clustering, though they differ in their network architectures and design strategies. For example, STAGATE first constructs a spatial neighbor network (SNN) based on spatial information, and then uses a graph attention autoencoder to learn low-dimensional latent embeddings by integrating both the SNN and gene expression data. SpaGCN is a graph convolutional network approach capable of integrating gene expression, spatial location and histology images.

The diversity of spatial domain detection methods, each with distinct assumptions, data requirements, strengths, and limitations, leads to varied outcomes, making it challenging to determine the most suitable approach for new spatial transcriptomics datasets ^[7,15,16]^. A commonly adopted approach to overcome this challenge involves designing ensemble frameworks that aggregate the outcomes of multiple individual algorithms for unsupervised clustering ^[17]^. These ensemble clustering approaches have been widely applied to both bulk and single-cell RNA-seq (scRNA-seq) data, such as COMSUC ^[18]^ for bulk multi-omics data, and SC3 ^[19]^, SAME-clustering ^[20]^, and GRMEC-SC ^[21]^ for scRNA-seq data. These methods have demonstrated significantly improved clustering accuracy and stability. However, to the best of our knowledge, there are very few published ensemble clustering approaches that integrate multiple spatial domain detection methods specifically designed for ST data, with the exception of EnSDD ^[22]^. EnSDD employs a weighted ensemble learning framework to construct a consensus similarity matrix by combining the results of multiple spatial domain detection methods, and subsequently applies the Louvain algorithm for spatial domain assignment. However, the weighted ensemble learning algorithms and most of the existing ensemble clustering methods face two main challenges: (i) ensemble methods aggregate information at the spot level but often fail to explore cluster-level connections; and (ii) ensembling focuses on the direct connection within the multiple base clustering, often overlooking the multiscale indirect relationships embedded in them.

To address the challenges associated with seamlessly integrating results from multiple spatial domain detection (SDD) methods (BayesSpace, BASS, STAGATE, SpaGCN, and SpatialPCA, which are representative methods selected from the two SDD categories) for ST data, we propose SpatialESD (**Spatial E**n**S**emble **D**omain detection), which leverages the enhanced ensemble clustering algorithm ^[23]^ to improve spatial domain detection. SpatialESD offers several advantages: (i) SpatialESD simultaneously detects spot-wise co-occurrence relationship as well as the multiscale indirect cluster-wise relationship between base spatial domain detection in ensembles; (ii) SpatialESD employs multiple advanced learners to obtain more stable, robust and accurate spatial domain detection for ST data than could be obtained from any spatial domain detection method alone.

We conducted simulation studies to evaluate the spatial domain detection performance of SpatialESD by comparing it with base clustering methods, as well as the EnSDD ensemble method based on the same base clustering methods. We further extensively evaluated SpatialESD on multiple human tissue datasets generated by 10x Visium, including annotated data of the human dorsolateral prefrontal cortex, breast cancer, and HER2 breast cancer, as well as unannotated data of invasive ductal carcinoma and ovarian cancer. Our results demonstrate that SpatialESD outperforms base methods and the EnSDD ensemble method based on the same set of base methods. With the domain detection results, we selected domain-specific differentially expressed genes and performed trajectory and cell-cell communication analyses, which provide valuable insights for developing novel cancer treatment strategies.

## 2 Methods

### 2.1 Overview of SpatialESD with the architecture analysis

SpatialESD leverages an enhanced ensemble clustering method to integrate spatial domain detection results derived from multiple SDD methods. To illustrate the idea, the current SpatialESD implementation embeds five such methods: BayesSpace, BASS, STAGATE, SpaGCN, and SpatialPCA. Users can also include other methods. An overview of the SpatialESD method is shown in Figure 1. The input data for SpatialESD includes gene expression profiles, spatial location, and histology images. The proposed SpatialESD consists of three steps (Figure 1a): (1) executing the five spatial clustering methods to identify spatial domains in the ST data; (2) constructing the enhanced co-association (ECA) matrix for the multiple spatial domain detection results; and (3) consensus detection of spatial domains using hierarchical clustering based on the ECA similarity matrix. In step (2), unlike the commonly used ensemble clustering strategy, which captures spot-wise similarity directly in ensemble clustering but fails to consider the potentially indirect relationships between base clusters, we first construct the Jaccard coefficients matrix as the initial similarity graph between clusters, and then enhance the cluster-wise similarity by incorporating multiscale direct and indirect relationships through a random walk process. Based on the multiscale cluster-wise similarity matrix, we further map it from the cluster level to the spot level, building the enhanced connectivity matrix for each base clustering. We then aggregate these enhanced connectivity matrices to form the final ECA for the entire ensemble. The ECA matrix effectively and efficiently captures multiscale direct and indirect relationships obtained from individual results, thus enhancing the final domain detection result.

**Figure 1.**
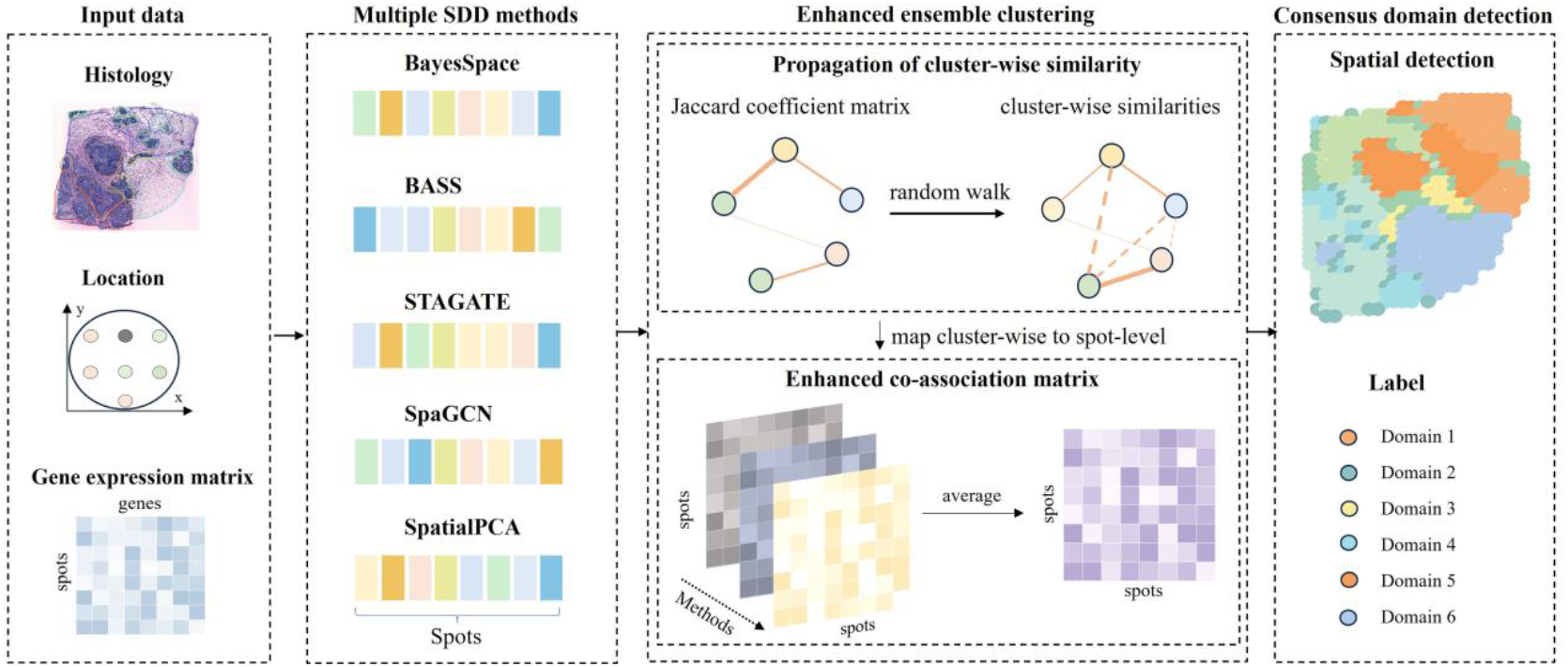
Workflow of SpatialESD. The input to SpatialESD consists of gene expression profiles, spatial coordinates, and histology images. SpatialESD first leverages the initial clustering results from multiple SDD methods to enhance cluster-wise similarities via the random walk algorithm. The cluster-wise similarity is then mapped to the spot-level to construct the ECA matrix. Finally, hierarchical clustering is applied to obtain consensus spatial domain detection.

Following the detection of spatial domain clusterings, subsequent downstream analyses are conducted, including differential gene detection, enrichment analysis, trajectory analysis, and cell-cell communication analysis, to reveal the biological characteristics and functions within different spatial domains, further advancing the understanding of complex biological systems.

### 2.2 Ensemble combination rule

Given ST data consisting of a gene expression matrix *X* ∈ ℝ^*N*×*p*^ and a spatial coordinates matrix *S* ∈ ℝ^*N*×2^, and corresponding histological image information *I*, where *N* is the number of spatial spots and *p* is the number of genes, we apply five spatial clustering methods, namely BayesSpace, BASS, SpaGCN, STAGATE and SpatialPCA, using their default parameter settings to obtain corresponding base clustering results. Each base clustering is denoted as 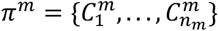, where *m* = 1, …, *M* and *n*_*m*_ is the number of clusters in the *m*-th method. In this work, *M* = 5. The set of base clustering is expressed as Π = {*π*^1^, …, *π*^*M*^}.

To facilitate the ensemble, we collect all cluster nodes from the selected base clustering into a unified set 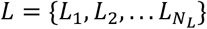, where *N*_*L*_ = *n*^1^ + … + *n* ^*M*^. *L*_*k*_ represents the *k*-th node, and *N*_*L*_ is the total number of nodes. These clusters are treated as nodes in a graph 𝒢 = {𝒱, ℰ}, where 𝒱 = *L* represents the set of nodes and ℰ represents the set of edges in the graph 𝒢, with edge weights between two nodes denoted as 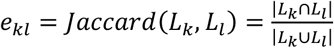. This graph provides the foundation for integrating multi-scale structural information in subsequent steps.

Unlike conventional co-association matrix methods that define spot similarity in a binary manner based solely on co-occurrence, e.g., assigning values of 0 or 1, SpatialESD introduces a similarity mapping mechanism. This allows the similarity between spots assigned to different clusters to be determined by the structural similarity between their respective clusters, thereby capturing finer-grained and more informative relationships. SpatialESD consists of three main steps: (1) Propagation of cluster-wise similarities, where multi-step random walks are performed on the cluster graph to integrate multi-scale structural information and capture the underlying multi-scale relationships between clusters; (2) Enhanced co-association matrix based on similarity mapping, which maps cluster-level similarities back to the spot-level to reflect the spot-level co-occurrence relationship as well as the cluster-wise structural information; and (3) Spatial domain consensus detection, obtained by performing hierarchical clustering on the ECA matrix.

#### Step 1: Propagation of cluster-wise similarities

To capture the structural relationships between clusters from different base clustering, we perform a random walk process on the cluster-wise graph to propagate multi-scale similarities. The one-step transition probability from node *L*_*k*_ to *L*_*l*_ is defined as: 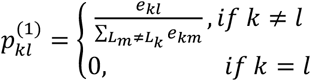.

From this initial transition probability matrix *P*^(1)^, we can further construct the *r*-step transition probability matrix *P*^(*r*)^ = *P*^(*r*−1)^ · *P, for r* > 1. For each node *L*_*k*_,we aggregate its random walk trajectories from step 1 to *r* into a trajectory vector 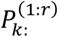.

Finally, we construct a cluster-wise similarity matrix 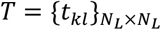 using the cosine similarity between the random walk trajectories:

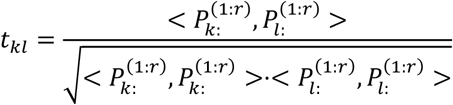

where <·,·> denotes the inner product. This matrix captures the multi-scale similarity structure between clusters and supports the downstream enhancement of spot-level relationships.

#### Step 2: Enhanced co-association matrix based on similarity mapping

After computing the cluster-wise similarity matrix *T*, we map the cluster-wise similarities back to the spot-level to build an ECA matrix 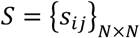, which incorporates both spot co-occurrence relationships and the multi-scale similarity information between clusters. The ECA matrix is obtained by averaging the enhanced connectivity matrices from the *M* base clusters: 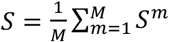. For the *m* -th base cluster, the enhanced connectivity matrix 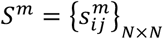 is defined as: 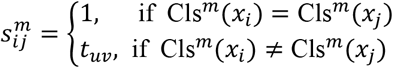. Where 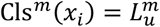 and 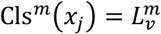, meaning that if spots *x*_*i*_ and *x*_*j*_ belong to the same cluster in the *m* -th base model, their similarity is set to 1. Otherwise, the similarity between nodes *u* and *v* is mapped to the similarity between spot *x*_*i*_ and *x*_*j*_, which is given by *t*_*uv*_.

#### Step 3: Spatial domain consensus detection

Given the final enhanced co-occurrence matrix *S*, we transform it into a distance matrix *D* = 1 − *S*, where higher similarity values in *S* correspond to smaller distances in *D*. Hierarchical clustering ^[24]^ with average linkage is then applied to *D* to obtain the final spatial domain consensus detection, which balances local pairwise relationships with global structural patterns.

### 2.3 Estimation of the number of clusters

Estimating the optimal number of clusters is critical for identifying spatial domains in unsupervised clustering. For datasets with manual annotations, the number of spatial clusters is set according to the ground truth. In the absence of such prior annotations, the optimal number of clusters *q* is determined by evaluating the average Negative Log-Likelihood (NLL) across different cluster numbers. Specifically, for each candidate cluster number *q*, we use a Markov Chain Monte Carlo (MCMC) algorithm to iteratively estimate model fit, implemented via the qTune function in the BayesSpace R package. To ensure model stability, the initial burn.in iterations (default 100) are discarded, and the average NLL is calculated from the remaining iterations. The negative log-likelihood, denoted as −log*L*(*q*), where *L*(*q*) is the log-likelihood given cluster number *q*. The optimal cluster number is chosen as the one that minimizes the average NLL. Here, *q* is set from 2 to 15 as the candidate range of cluster numbers, and we set the total number of iterations as 1000. We use the qPlot function to show the relationship between cluster number and NLL, and select the value with the lowest or most stable NLL as the final cluster number, which typically corresponds to the “elbow” point in the plot.

### 2.4 Base spatial domain detection algorithms

In this study, we selected three statistical methods BayesSpace, BASS, and SpatialPCA, and two graph-based deep learning methods SpaGCN and STAGATE as the base spatial domain detection algorithms. A brief overview of the characteristics of the base clustering method is provided in Table S1. We compared the spatial domain detection performance of SpatialESD with base clustering methods, as well as the EnSDD ensemble method ^[22]^ based on the same base clustering to ensure comparability.

BayesSpace ^[9]^ is a Bayesian spatial clustering method for spatial domain detection in spatial transcriptomics data generated from the Visium or ST platforms. BayesSpace performs principal component analysis (PCA) on the normalized gene expression matrix and models the top 15 principal components (PCs) as a low-dimensional representation. In this study, we followed the original BayesSpace work and used the default parameters. Specifically, we selected the top 2,000 highly variable genes (HVGs) which were used to compute the top 15 PCs as input features for clustering.

BASS ^[10]^ is a multi-scale and multi-sample analytical method designed for spatial transcriptomics, enabling simultaneous cell type clustering and spatial domain detection within a unified modeling framework. It employs a Bayesian hierarchical model that jointly incorporates normalized gene expression data and spatial location information. Principal components obtained via PCA on the gene expression matrix are used as model inputs. BASS treats the cell type and spatial domain labels for each cell as latent variables and infers them using an efficient sampling algorithm. The method also automatically estimates the spatial interaction parameter *β*, allowing it to effectively capture spatial dependencies among neighboring cells. In this study, we followed the original implementation and default parameter settings of BASS. For the DLPFC dataset, which contains multiple tissue sections, we applied BASS in its multi-sample analysis mode to improve the accuracy of spatial domain detection. For all other datasets, we performed single-sample analyses by running the model independently on each individual tissue section.

STAGATE ^[13]^ is an unsupervised learning method based on a Graph Attention Auto-Encoder, designed to identify spatial domains in spatial transcriptomics data by integrating gene expression profiles and spatial location information. The method constructs a spatial neighbor graph to represent the spatial relationships among spots. STAGATE employs an attention mechanism to adaptively learn the similarity between neighboring spots, generating low-dimensional latent representations that capture spatial expression patterns. To enhance the detection of domain boundaries, STAGATE incorporates an attention mechanism for more accurate modeling of spatial similarity. In this study, we followed the original implementation of STAGATE and applied its default parameter settings to analyze the spatial transcriptomics data.

SpaGCN ^[4]^ is a graph convolutional network (GCN)-based method designed to identify spatial domains and detect spatially variable genes (SVGs) by integrating gene expression, spatial location, and histology information in spatial transcriptomics data. The method first performs principal component analysis (PCA) on the normalized gene expression matrix and uses the top 50 principal components as the low-dimensional representation. SpaGCN then constructs an undirected weighted graph that incorporates spatial proximity and histological similarity among spots. A graph convolutional layer is applied to aggregate gene expression information from neighboring spots, followed by an unsupervised iterative clustering algorithm to identify spatial domains. In this study, we adopted the original implementation and default parameter settings of SpaGCN. For datasets with histology information, SpaGCN incorporated histology-derived features into the graph construction process. For datasets lacking histological images, only spatial coordinates were used to compute distances between spots, and histology information was excluded from the spatial graph construction.

SpatialPCA ^[12]^ implements a probabilistic principal component model for spatial dimensionality reduction designed to identify spatial domains in spatial transcriptomics data by integrating gene expression profiles and spatial location information. SpatialPCA leverages a spatial correlation kernel matrix to perform dimensionality reduction on gene expression data, generating low-dimensional latent representations that explicitly preserve spatial correlation patterns across tissue locations. To enhance the detection of domain boundaries, SpatialPCA further implements spatial smoothing for more accurate modeling of spatial similarity. In this study, we applied the default parameter settings to analyze the spatial transcriptomics data.

EnSDD ^[22]^ is an unsupervised ensemble learning framework designed to improve the accuracy and robustness of spatial domain detection by integrating multiple clustering results. For each base clustering, a binary similarity matrix is constructed to indicate whether any two spots belong to the same cluster. EnSDD jointly learns a consensus similarity matrix and a set of weights for the base clusters by minimizing the weighted Euclidean distance between each base similarity matrix and the consensus, with an additional negative entropy regularization term to avoid dominance by any single clustering. Once the consensus matrix is obtained, the Louvain algorithm with an adaptive resolution parameter is applied to generate the final spatial domain assignments. In this study, to ensure a fair comparison between EnSDD and our proposed method, we used the same base clustering results as input for both methods. This allowed the ensemble step in EnSDD to operate on the same underlying clustering information, ensuring consistency and comparability between the two approaches.

### 2.5 Evaluation metrics

#### ARI

We utilize the ARI ^[25]^ to assess the spatial clustering performance of different methods. ARI is an improved variant of the Rand index (RI), used to measure the agreement between two clustering results. Its value ranges from -1 to 1, with higher scores indicating stronger consistency between partitions. We calculate the ARI value using the adjustedRandIndex function within the R package mclust.

#### LISI

Local inverse Simpson’s index (LISI) ^[26]^ is used to quantify the clustering performance for spatial domain detection. The LISI score reflects the effective number of spatial domain labels present in the local spatial neighborhood of a given spot, and it is calculated as:

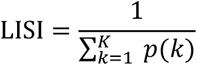

where *p*(*k*) is the probability of the spatial domain cluster *k* appearing in the local neighborhood, and *K* is the total number of spatial domains. The LISI score is computed using the “compute_lisi” function from the LISI R package with the default parameter (perplexity=15). A lower LISI score indicates that the spot’s local neighborhood is composed of more homogeneous spatial domain clusters, reflecting better spatial clustering performance.

#### PAS

PAS score ^[12]^ evaluates the spatial inconsistency of spots assigned to a cluster. It is defined as the proportion of a spot’s *k* nearest neighbors (*k*=10) that have a different cluster label. A spot is considered spatially inconsistent when 60% or more of its neighbors belong to other clusters. A lower PAS score indicates better spatial domain clustering, reflecting greater homogeneity within spatial clusters. In other words, a smaller PAS score suggests that the spots within a given cluster are more spatially coherent and less likely to be randomly distributed.

## 3 Simulation study

### 3.1 Simulation design

We performed comprehensive and realistic simulations to evaluate the performance of SpatialESD and compared it with its corresponding base clustering methods, as well as with the EnSDD ensemble method applied to the same set of base clustering. We leveraged the SRTsim ^[27]^ framework to generate synthetic datasets that preserve both gene expression characteristics and spatial patterns of the reference dorsolateral prefrontal cortex (DLPFC) ^[28]^ tissue slice (sample 151670, containing 3,498 spatial spots, 18,311 genes, and 5 annotated tissue domains). The simulation process followed a tissue-based strategy, wherein gene expression profiles and spatial coordinates from the reference dataset were utilized to ensure structural and transcriptional fidelity. Spatial coordinates from the original dataset were directly incorporated into the synthetic data to maintain the spatial architecture of the tissue.

For each gene in the reference dataset, four count-based models, namely Poisson, zero-inflated Poisson (ZIP), negative binomial (NB), and zero-inflated negative binomial (ZINB), were fitted. The optimal model was selected based on the minimum Akaike Information Criterion (AIC), and its parameters were employed to simulate gene-specific expression counts, preserving inherent features such as overdispersion and zero inflation. To retain spatial expression patterns observed in the reference data, SRTsim implemented a rank-ordering allocation mechanism. This involved ranking spatial spots according to gene expression intensity in the reference dataset and assigning simulated counts to corresponding spots in the synthetic dataset following the same rank order. This approach ensured that spatial gradients and domain-specific expression patterns were meticulously reproduced.

To validate simulation robustness, the process was repeated 20 times with distinct random seeds. Outputs included spatial coordinate matrices, gene expression count matrices, estimated model parameters, and random seeds, ensuring full reproducibility of the synthetic datasets. The accuracy of spatial domain detection in simulated data was quantified based on ARI, which measured concordance between clustering results and ground-truth domain labels. Additionally, six evaluation metrics were applied to assess global data fidelity: gene-level metrics (mean expression, variance, coefficient of variation, and zero proportion) and spot-level metrics (zero proportion and library size). These analyses collectively validated the ability of SRTsim to generate realistic synthetic ST data that recapitulate both transcriptional and spatial features of the original reference.

### 3.2 Simulation results

We evaluated the performance of SpatialESD using 20 simulated datasets generated based on sample 151570 from the 10x Visium DLPFC sample. By comparing the synthetic data with the reference data, we evaluated how well the simulated data retained the characteristics of the original data. The results (Figure 2a) show that, across these six metrics, the simulated data closely resemble the reference data. This indicates that the synthetic data effectively preserves the key biological and technical features of the original data, validating the ability of the simulation methods to reproduce the characteristics of real data. The median ARI value from the 20 simulations for each algorithm’s clustering result was taken as the final evaluation result. The experimental results are shown in Figure 2b. The ARI value of SpatialESD is overall higher than that of other base spatial domain detection methods, namely BayesSpace, BASS, SpaGCN, STAGATE, and SpatialPCA, as well as the EnSDD ensemble strategy, demonstrating the superior accuracy of SpatialESD in spatial domain detection. The detailed ARI results are presented in Table S2. Furthermore, the visualization of spatial domain detection (Figure 2c) shows that the SpatialESD method more clearly and accurately identifies the layer_3 region. It also outperforms STAGATE and SpatialPCA methods in recognizing the WM layer.

**Figure 2.**
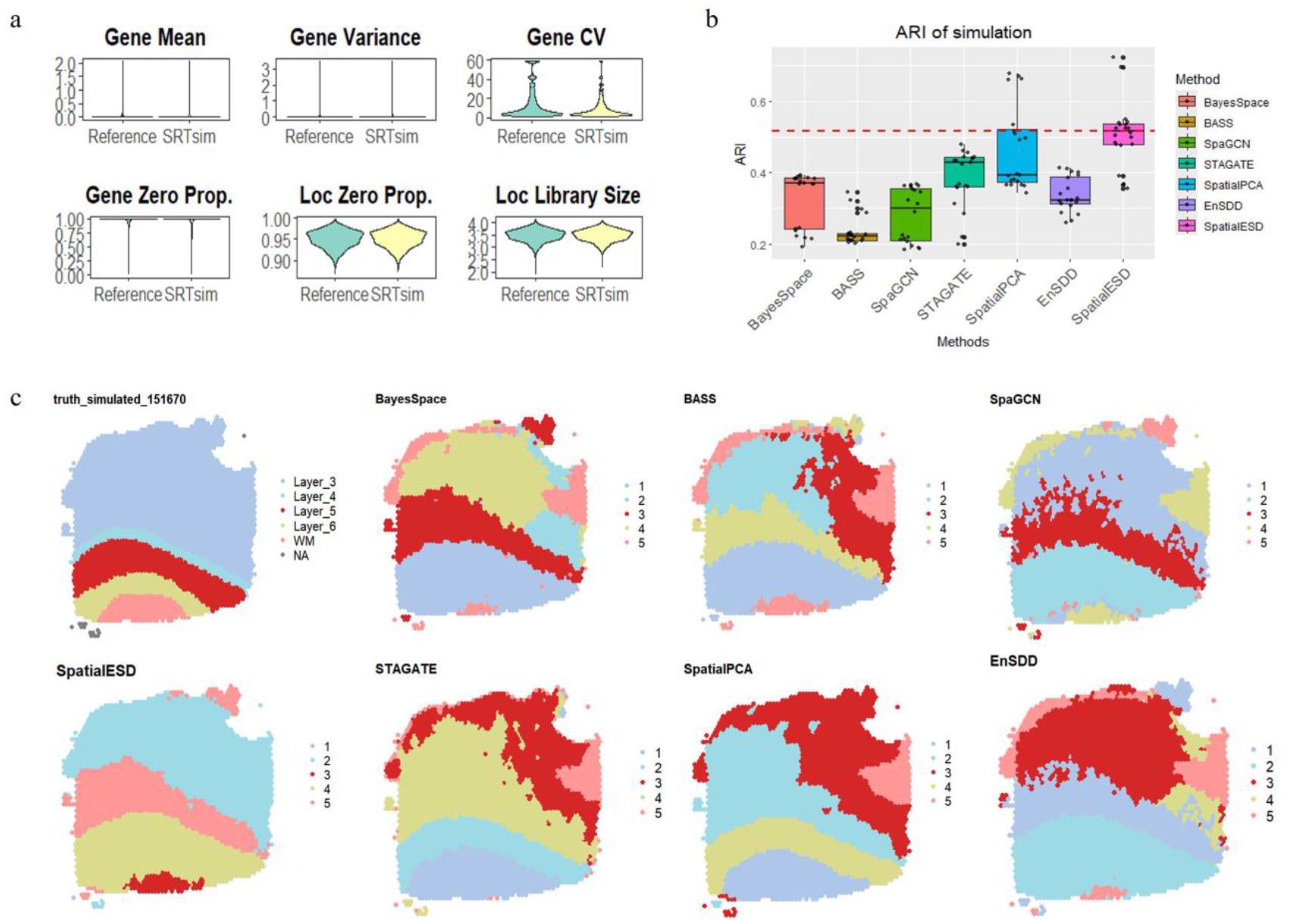
SpatialESD enhances tissue structure detection in simulated data. (a) The violin plots show the distributions of six metrics in the reference and synthetic datasets. (b) Boxplots of the 20 ARI values for the six methods. (c) Visualization of domain detection results, including the ground truth and those by different methods.

## 4 Real data analysis

### 4.1 Data analyzed

#### Human DLPFC ST data

The human dorsolateral prefrontal cortex (DLPFC) ST data, generated using the 10x Visium platform, is widely used as a benchmark for evaluating spatial domain detection methods. It includes 12 postmortem tissue sections from three neurotypical adult donors, with each section containing 3,460 to 4,789 spatial spots and expression profiles for 33,538 genes. Based on histological features, manual annotations define five to seven distinct regions, including cortical layers (Layers 1–6) and white matter (WM) ^[28]^. These annotations provide ground truth labels for performance assessment. The processed dataset is publicly accessible via spatialLIBD (https://github.com/LieberInstitute/spatialLIBD).

#### Human breast cancer ST data

The human breast cancer ST data were generated using the 10x Visium platform, consisting of 3,798 spots and expression data for 36,601 genes ^[29]^. This dataset represents complex microenvironments and highly heterogeneous characteristics of breast cancer. The data have been manually annotated by a pathologist based o n H&E images and the spatial expression profiles of known breast cancer marker genes. This annotation serves as the ground truth in our application and is available at https://github.com/JinmiaoChenLab/SEDR_analyses/blob/master/data/BRCA1/metadata.tsv.

#### Human HER2-positive breast cancer ST data

ST data for HER2-positive breast cancer were generated using the 10x Visium platform. This dataset includes spatial gene expression and cell type information from 36 samples collected from 8 HER2-positive individuals (patients A-H). Each tumor (*n* = 36 slices) had 3 (adjacent) or 6 (evenly distributed) slices ^[30]^. Each slice contains between 176 to 691 spots, capturing expression data for approximately fifteen thousand genes. A pathologist examined and annotated one slice of each tumor based on morphology in the corresponding HE image (hematoxylin and eosin). Regions were labeled as carcinoma in situ, invasive carcinoma, adipose tissue, immune infiltration, or connective tissue. The data is available at https://github.com/almaan/her2st.

#### Human ovarian cancer ST data

ST data for human ovarian cancer data were generated using the 10x Visium platform, including 3,493 spots and 36,601 genes. Zhao et al. ^[9]^ performed immunofluorescence staining on an ovarian cancer (OC) sample (1 tissue slice, *n* = 3,493 spots) and conducted cell segmentation based on the IF staining. This segmentation data serves as a reference in our analysis.

#### Human invasive ductal carcinoma (IDC) ST data

The human IDC ST data were generated using the 10x Visium platform, including 4,727 spots and 36,601 genes. Zhao et al. ^[9]^ performed immunofluorescence staining on an IDC sample (1 tissue slice, *n* = 4,727 spots) and conducted cell segmentation based on histological annotations. Regions were labeled as invasive carcinoma, carcinoma in situ, benign hyperplasia, unclassified tumor, and non-tumor areas.

### 4.2 Downstream analysis

#### 4.2.1 Identifying spatial domain-specific DEGs

SpatialESD provides two methods to identify genes with domain-specific spatial expression patterns, especially those showing significant differences across different spatial domains. The first approach uses the Wilcoxon test ^[31]^ to identify domain-specific differentially expressed genes (DEGs), focusing on genes with a log2 fold change greater than or equal to 1. These genes exhibit differences in expression levels between the selected domain and other spatial domains.

The second approach leverages spatial autocorrelation (SA) analysis to characterize the spatial dependency of gene expression across tissue locations ^[32]^. SA measures the extent to which gene expression values at nearby spots are correlated. To further assess whether these DEGs exhibit localized spatial clusters within specific domains, we calculate the Local Getis and Ord’s *G*_*i*_ ^[33]^, which quantifies the degree of local spatial clustering of gene expression at each spot. This metric ranges from negative to positive values, where higher positive values indicate stronger local spatial aggregation, suggesting the presence of “hot spots” of elevated expression. Mathematically, the Local Getis and Ord’s *G*_*i*_ is defined as:

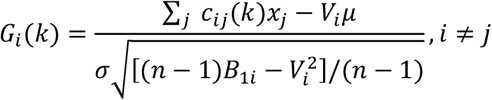

where *x*_*j*_ represents the gene expression in spot *j, μ* and *σ* are the mean and standard deviation of gene expression across all spots, excluding spot *i. c*_*ij*_(*k*) denotes the spatial distance between spots *i* and *j*, and *k* is the number of spots closest to spot *i. V*_*i*_ = ∑_*j*_ *c*_*ij*_ and 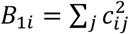.

Local Getis and Ord’s *G*_*i*_ statistic is robust to the choice of *k* which is set as 6 in our analysis. We use the localG function from the R package spdep for the calculation of Local Getis and Ord’s *G*_*i*_ statistic.

#### 4.2.2 Enrichment analysis

We performed functional enrichment analysis on the differentially expressed genes identified from differential expression analysis using the KOBAS 3.0 online platform ^[34]^. Specifically, Gene Ontology (GO) ^[35]^ annotation and Kyoto Encyclopedia of Genes and Genomes (KEGG) ^[36]^ pathway enrichment analyses were conducted to identify biological processes and signaling pathways potentially associated with disease progression. Significance thresholds were defined as both *P*-value < 0.01 and FDR-adjusted *P*-value < 0.01. Enriched pathways were visualized using bubble plots to illustrate the proportion of genes involved in each pathway.

#### 4.2.3 Trajectory analysis

We applied Slingshot ^[37]^ on the low-dimensional components to uncover potential developmental trajectories among spatial locations within the tissue. Cluster labels obtained from SpatialESD were used as input to guide the inference. The Slingshot algorithm was employed to construct trajectories and assign a pseudo-time value to each spatial location. As is common in most lineage inference methods, it was necessary to designate a starting point, typically a specific cluster or cell, based on biological knowledge. While this choice defines the direction of the trajectory, it does not affect the relative positions of clusters along it. For the IDC data, we specified the non-tumor region as the start cluster, and for the OC data, the trajectory was anchored at the CD45 region.

#### 4.2.4 Cell communication analysis

We performed cell-cell communication analysis using CellChat v2 ^[38]^, which includes an expanded ligand-receptor database, additional functional annotations, and new comparison features. A gene expression matrix normalized by the library size and log-transformation was used as input. Cells were annotated with corresponding cell type labels (e.g., tumor cells, immune cells, fibroblasts), based on spot-level predictions using Seurat and reference single-cell data from HumanPrimaryCellAtlasData ^[39]^, following the annotation procedure described in the Seurat spatial vignette (https://satijalab.org/seurat/articles/spatial_vignette.html). We then constructed a CellChat object using the expression matrix and the annotated cell type labels, referencing CellChatDB.human for ligand-receptor interactions and signaling pathways. Communication probabilities between cell types were calculated based on the average expression of ligands and receptors using the truncated mean method. The resulting interaction networks were visualized using netVisual_aggregate, netVisual_individual, and netVisual_diffInteraction, while the roles of each cell group in sending or receiving signals were assessed via the function netAnalysis_signalingRole_scatter and netAnalysis_signalingRole_heatmap. These analyses enabled the detection of dominant signaling cell types and active pathways involved in intercellular communication.

### 4.3 Results

#### 4.3.1 Application to the DLPFC ST data

The 12 DLPFC slices were analyzed using ensemble clustering with five base algorithms. The detailed ARI values were presented in Table S3. The boxplot (Figure 3a) demonstrates that SpatialESD achieved the most superior overall performance on this dataset, attaining the highest accuracy and exhibiting the most robust performance. For spatial domain visualization analysis, we conducted an in-depth examination of slice 151671, where the ARI value increased from a minimum of 0.504 in base clustering to 0.833 with SpatialESD. Visualization results (Figures 3b-c) reveal that some base clustering methods, such as STAGATE and SpatialPCA, struggled to accurately identify the layer_3 region in the slice, and nearly all methods failed to detect the layer_4 region, while SpaGCN showed confusion in delineating the boundaries of the layer_6 region. In contrast, SpatialESD’s spatial domain visualization results align more consistently and precisely with manual annotations, fully identifying the layer_3 region and displaying clear boundaries for the layer_6 region. Although SpatialESD, like all other methods, exhibited limitations in distinguishing the layer_4 region and showed slightly vague boundary delineation, it still demonstrated superior overall performance compared to the alternatives.

**Figure 3.**
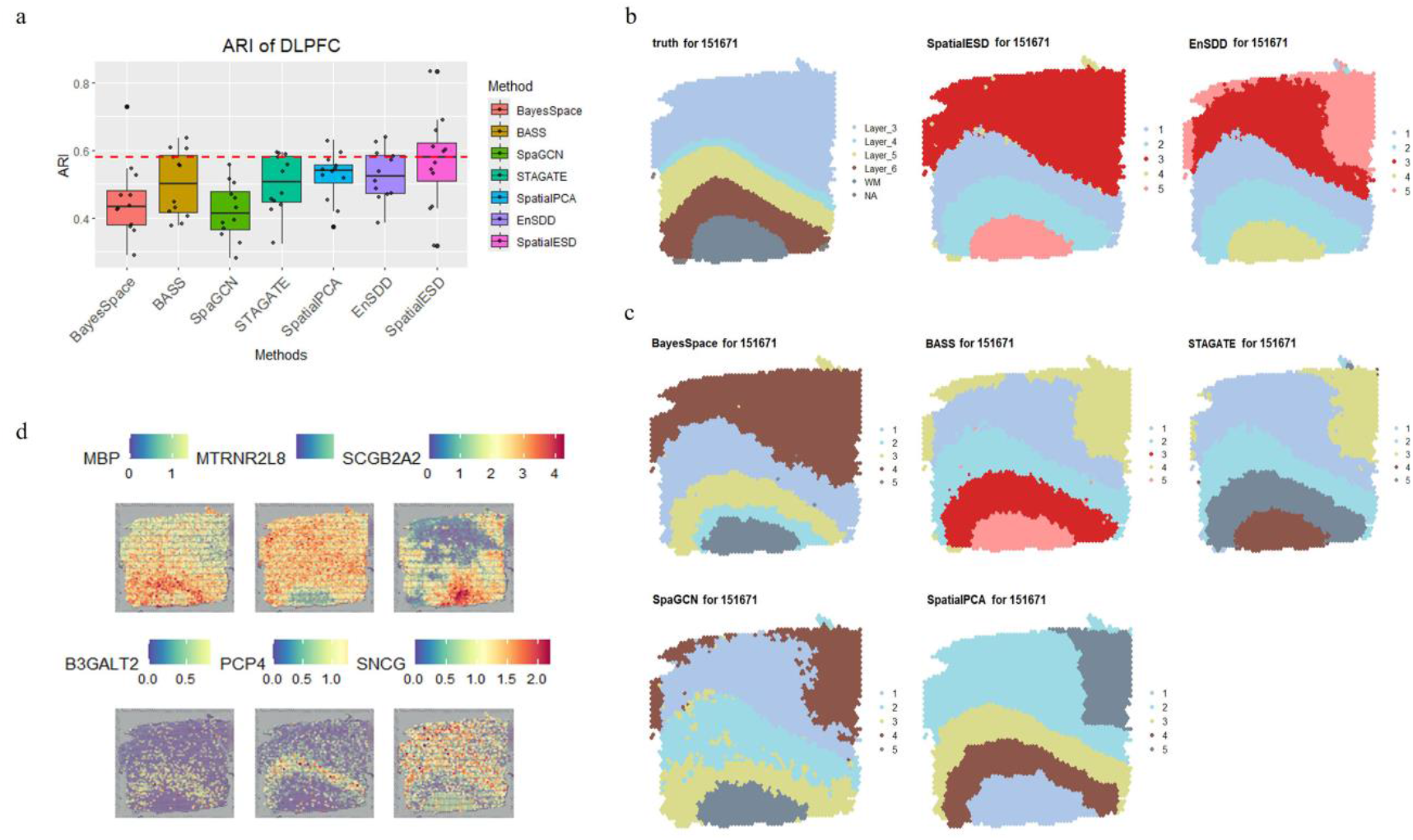
SpatialESD enhances tissue structure detection in DLPFC. (a) Boxplots of ARI values for the 12 tissue slices. (b-c) Visualization of spatial domains using the SpatialESD method and other methods. (d) Visualization of differentially expressed genes selected based on the Wilcoxon test.

Based on the spatial domains identified by SpatialESD, we first examined the spatial expression patterns of representative genes. As shown in Figure 3d, genes such as *MBP* and *PCP4* exhibited distinct region-specific expression across the tissue section. *MBP* exhibited high expression in the WM region, which is consistent with previously reported findings ^[28]^. *PCP4*, mainly expressed in Layer_5, is a calmodulin-binding protein implicated in neuronal signaling and plasticity ^[40]^. These spatially resolved expression patterns highlight the functional and compositional heterogeneity across different tissue regions. Full results of the downstream analysis for DLPFC are provided in Figure S1-Figure S5.

#### 4.3.2 Application to the breast cancer ST data

The Breast cancer dataset involved tissue images segmented into 20 regions and classified into four major morphological types, including ductal carcinoma in situ/lobular carcinoma in situ (DCIS/LCIS), invasive ductal carcinoma (IDC), healthy tissue, and tumor edge with low malignant features, manually annotated by Xu et al. ^[29]^. We selected four base clustering methods, BayesSpace, BASS, SpaGCN, and STAGATE, to ensemble, considering the low accuracy of SpatialPCA. To ensure reliability, we conducted ten repeated analyses by varying the random seed and assessed the clustering accuracy of SpatialESD and the base methods, with the detailed ARI results provided in Table S4 in the supplemental file. The boxplot (Figure 4a) demonstrates that SpatialESD achieved the most superior overall performance on the breast cancer dataset, attaining the highest accuracy and exhibiting the most robust performance. In the spatial domain visualization analysis (Figures 4b-c), BayesSpace exhibited significant confusion in identifying the IDC_6 region, SpaGCN and BayesSpace produced blurred delineations of the Healthy1 region, SpatialPCA performed chaotically in the IDC_4 region, and all base clustering methods performed poorly in the IDC_2 region. In contrast, the visualization results of SpatialESD across all regions exhibited higher consistency with the manual annotations, consistently outperforming other individual spatial domain detection methods. Full details of the downstream analyses are provided in the supplementary figures (Figure S6-Figure S10).

**Figure 4.**
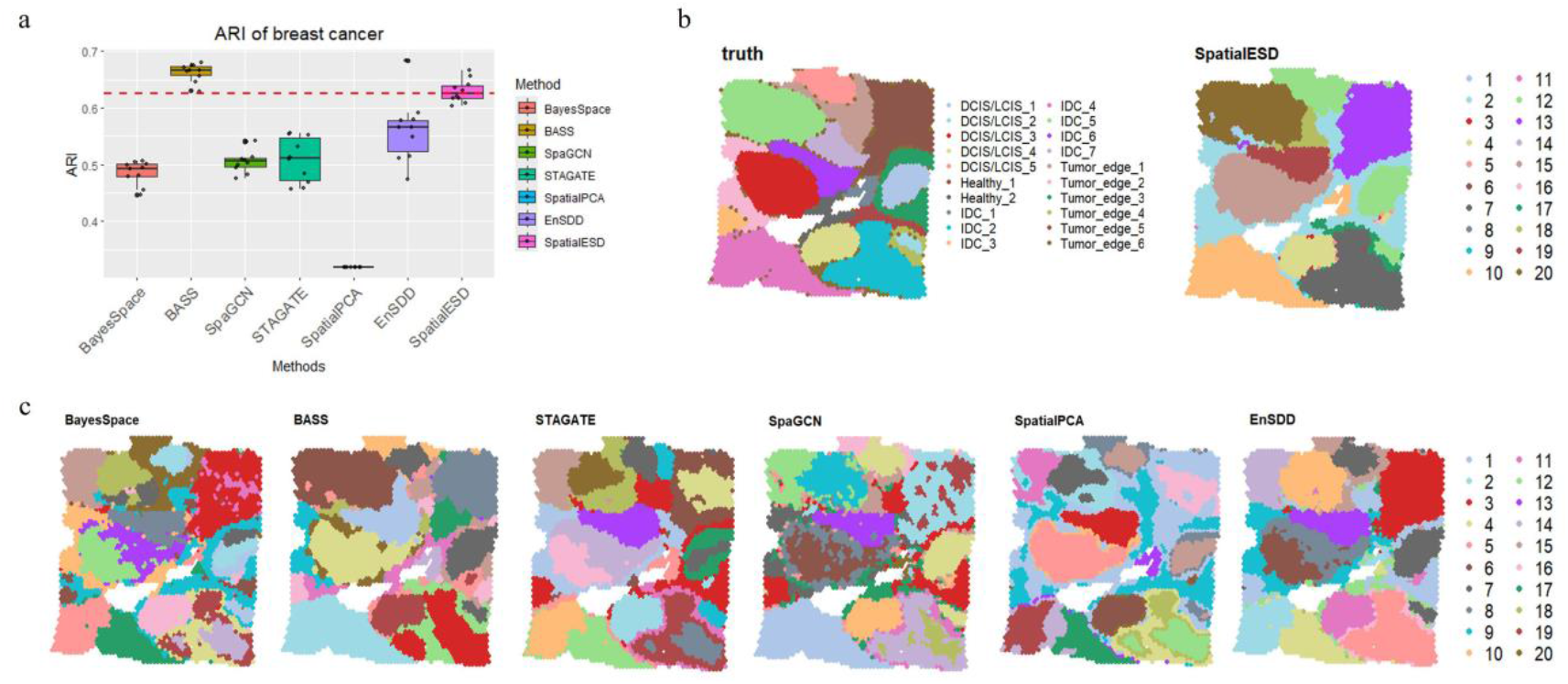
SpatialESD enhances tissue structure detection in breast cancer. (a) Boxplots of ARI. (b-c) Visualization of spatial domains using the SpatialESD ensemble method and several SDD methods.

#### 4.3.3 Application to the HER2 ST data

We also applied SpatialESD to an eight-slice HER2 dataset manually annotated by Andersson ^[30]^. The results show that SpatialESD achieved a slightly lower ARI value than BASS (by 0.016) but still outperformed other methods, exceeding them by 0.041 to 0.106, and further demonstrate the superior visualization compared to both individual spatial methods and EnSDD (Supplementary Note 1).

#### 4.3.4 Application to the Invasive Ductal Carcinoma ST data

We also analyzed the Invasive Ductal Carcinoma ST data which is unannotated. Using an elbow plot (Figure 5a), we determined the optimal number of clusters for the IDC dataset to be 10 and repeated the clustering results 10 times with varying seed numbers to compute the median ARI as an evaluation metric. Moreover, we also considered an alternative scenario where we repeated the clustering results for cluster numbers ranging from 2 to 10. We evaluated the clustering results using unsupervised clustering metrics, including the LISI and PAS metrics. The consistency of their trends with ARI on the DLPFC dataset (Figure 5b) supports the validity of these metrics as reliable evaluation tools. The results (Figure 5c) show that SpatialESD achieves superior performance compared to both base SDD methods and EnSDD ensemble approaches, with notable improvements in clustering accuracy and stability under both IDC dataset scenarios. The detailed ARI results are presented in Supplementary Tables S6-S9. The visual results (Figure 5d) show that the SpatialESD method more accurately distinguishes the invasive and in situ regions. In contrast, other clustering methods perform poorly, particularly in delineating the contours of the in situ region and in distinguishing non-tumor areas, where the boundaries are often confused. This further demonstrates that SpatialESD outperforms other methods in terms of overall performance.

**Figure 5.**
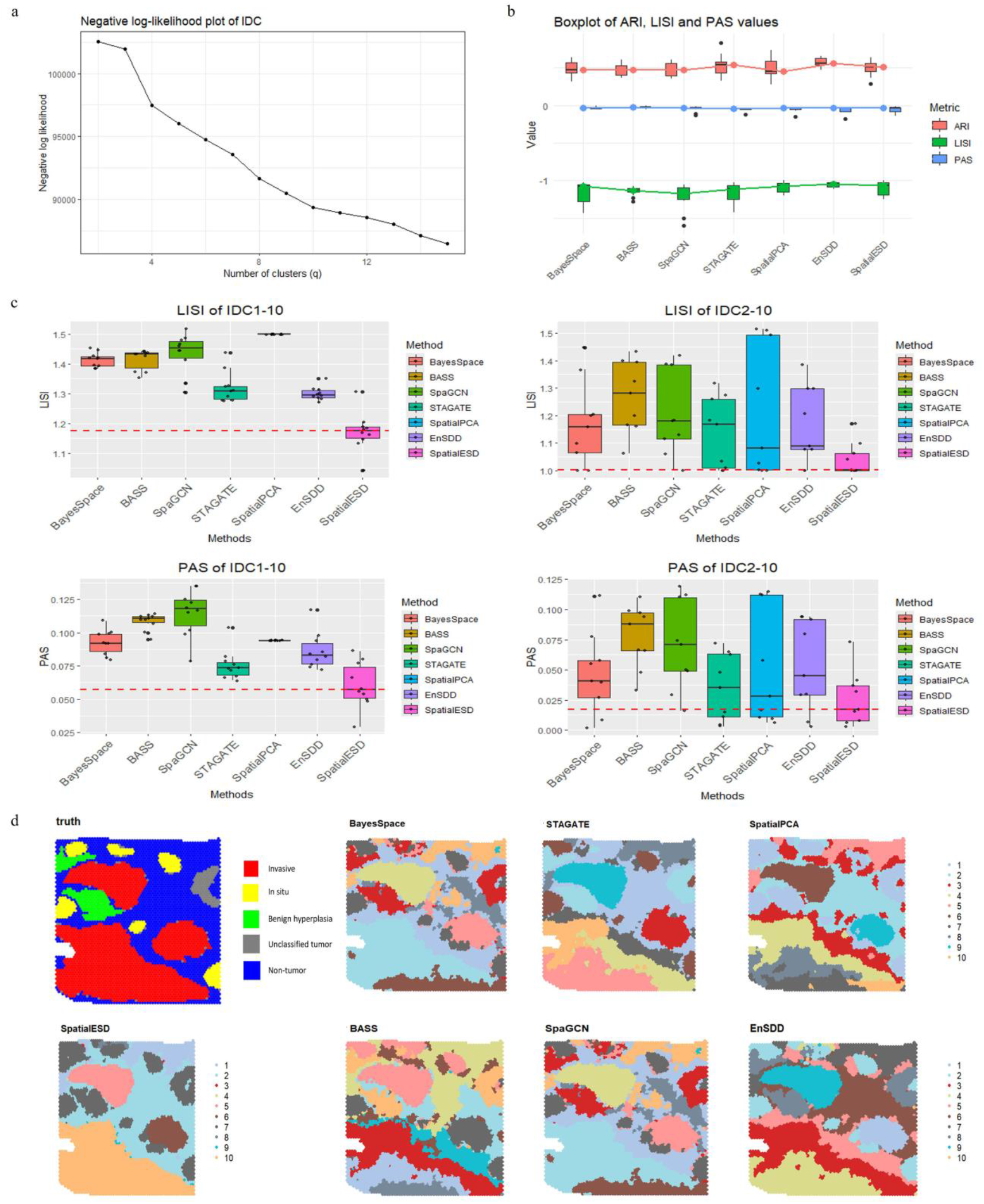
SpatialESD enhances tissue structure detection in invasive ductal carcinoma. (a) The optimal number of clusters for the IDC dataset is determined to be 10 based on the elbow plot. (b) Trend plots for annotated metric ARI and unannotated metrics LISI and PAS. (c) Boxplot results for the two evaluation metrics in the IDC dataset under two different scenarios. (d) Spatial domain visualizations of different methods.

Based on the spatial domains detected by SpatialESD, we employed two different methods, the Wilcoxon test (Figure 6a) and spatial autocorrelation (Figure 6b), to detect differentially expressed genes specific to each spatial domain. Figure 6a shows the visualization of differentially expressed genes in regions such as invasive, non-tumor areas, and in situ. We identified genes associated with a specific spatial domain type through differential gene analysis. Among these, genes such as *SLC39A6, KRT37, MGP, IFI27, BAMBI*, and *CXCL14* were selected from other regions, which are related to the biological characteristics and microenvironment of the tumor. *SLC39A6* has been identified as a key gene involved in regulating estrogen and epithelial-mesenchymal transition (EMT), and its expression is associated with poor survival outcomes in cancer patients ^[41]^. *KRT37* is involved in the estrogen signaling pathway and has been associated with cancer-related pathways, showing differential expression patterns that may influence cancer progression and treatment response ^[42]^. *MGP* has been identified as a potential breast-specific marker, with high sensitivity in detecting various subtypes of breast carcinoma, including triple-negative breast cancer and metaplastic carcinoma ^[43]^. *IFI27* is highly expressed in several cancers, including pancreatic cancer, where its high expression levels are associated with poorer overall survival and correlated with changes in immune cell infiltration, such as a decrease in CD8+ T cells and an increase in M2 macrophages ^[44]^. *BAMBI* is involved in regulating stemness and promoting the growth and maintenance of cancer stem cells, influencing tumor growth and metastasis ^[45]^. *CXCL14* plays a role in tumor metastasis, being predominantly expressed in fibroblasts, and influences cancer progression through multiple mechanisms ^[46]^. Figure 6b displays the Local Getis and Ord’s *G*_*i*_ values for the top six genes, differentially expressed in the first spatial domain compared to other domains, across all categories as detected by SpatialESD. The results indicate that the *G*_*i*_ values for these six genes in the first spatial domain are generally higher than in other categories, suggesting that these genes exhibit similar expression patterns at neighboring spatial points, demonstrating strong spatial autocorrelation.

**Figure 6.**
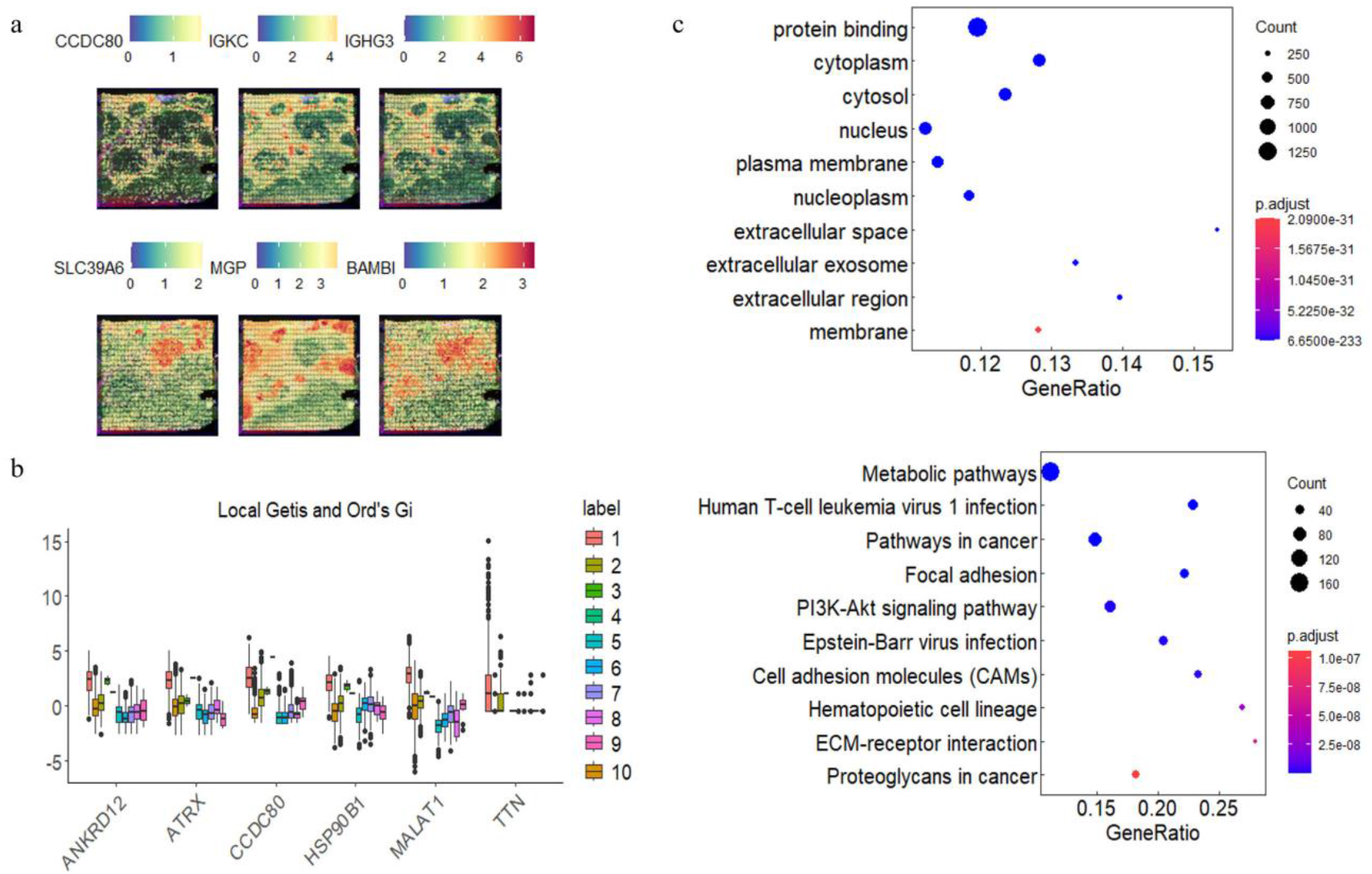
The downstream analysis corresponding to the IDC dataset. (a) Visualization of differentially expressed genes selected using the Wilcoxon test. (b) Expression levels of differentially expressed genes in different spatial domains based on spatial autocorrelation analysis. (c) Enrichment analysis of differentially expressed genes (upper: GO enrichment; lower: KEGG pathway enrichment).

Genes with both *P*-value≤0.05 and adjusted *P*-value≤0.05 were selected, resulting in 2,241 differentially expressed genes. These genes were then subjected to functional enrichment analysis, including the GO and KEGG pathways analysis. The GO analysis (Figure 6c, upper panel) revealed that the top ten functional enrichments are mainly associated with biological processes such as protein binding, cytoplasm, cytosol, nucleus, plasma membrane, nucleoplasm, various extracellular components, extracellular space, extracellular exosome, and extracellular region. In particular, the enrichment in membrane-bound and extracellular compartments points to potential involvement in intercellular communication and molecular transport. These cellular and molecular characteristics are critical in the context of tumor progression, influencing cell signaling, tumor-stroma interactions, and modulation of the tumor microenvironment ^[47-49]^. The biological pathways showed (Figure 6c, lower panel) a strong association with pathways involved in cancer-related signaling, such as Metabolic pathways, Human T-cell leukemia virus 1 infection, Pathways in cancer, Focal adhesion, PI3K-Akt signaling pathway, Epstein-Barr virus infection, Cell adhesion molecules (CAMs), Hematopoietic cell lineage, ECM-receptor interaction, and Proteoglycans in cancer. The enriched pathways are potentially involved in various biological processes and regulatory mechanisms related to cancer progression and the tumor microenvironment. The ECM-receptor interaction pathway regulates the interaction between tumor cells and the extracellular matrix, promoting tumor invasion, metastasis, and proliferation ^[50]^. Human T-cell leukemia virus 1 infects CD4 T lymphocytes and disrupts immune signaling networks, promoting the survival and dissemination of infected cells, thereby triggering the development of cancer in the host ^[51]^. The PI3K/Akt signaling pathway is involved in differential prognosis, and its activity is significantly reduced when DLAT is knocked down ^[52]^. Epstein-Barr virus (EBV) infection was detected in breast invasive ductal carcinoma and was significantly associated with nodal involvement ^[53]^. The complete results of the downstream analysis are provided in Supplementary Note 2.

#### 4.3.5 Application to the Ovarian Cancer ST data

We further analyzed the ovarian cancer (OC) dataset which has no manual annotations. The optimal cluster number was determined as 8 using an elbow plot (Figure S17), and the clustering process was repeated 10 times by varying the seed number, with the median ARI calculated as the evaluation metric. Additionally, similar to the IDC dataset, we explored another scenario where the cluster number was set from 2 to 10, followed by multiple repeated clustering analyses. As shown in Figure 7a, SpatialESD consistently outperformed both base SDD methods and EnSDD ensemble strategy across both scenarios, exhibiting higher clustering accuracy and greater stability. The detailed ARI results are presented in Supplementary Tables S10-S13. Visualization results (Figure 7b), when compared with H&E staining outcomes, reveal that SpatialESD achieved superior accuracy in identifying the CD45 region, while other clustering methods exhibited poor performance.

**Figure 7.**
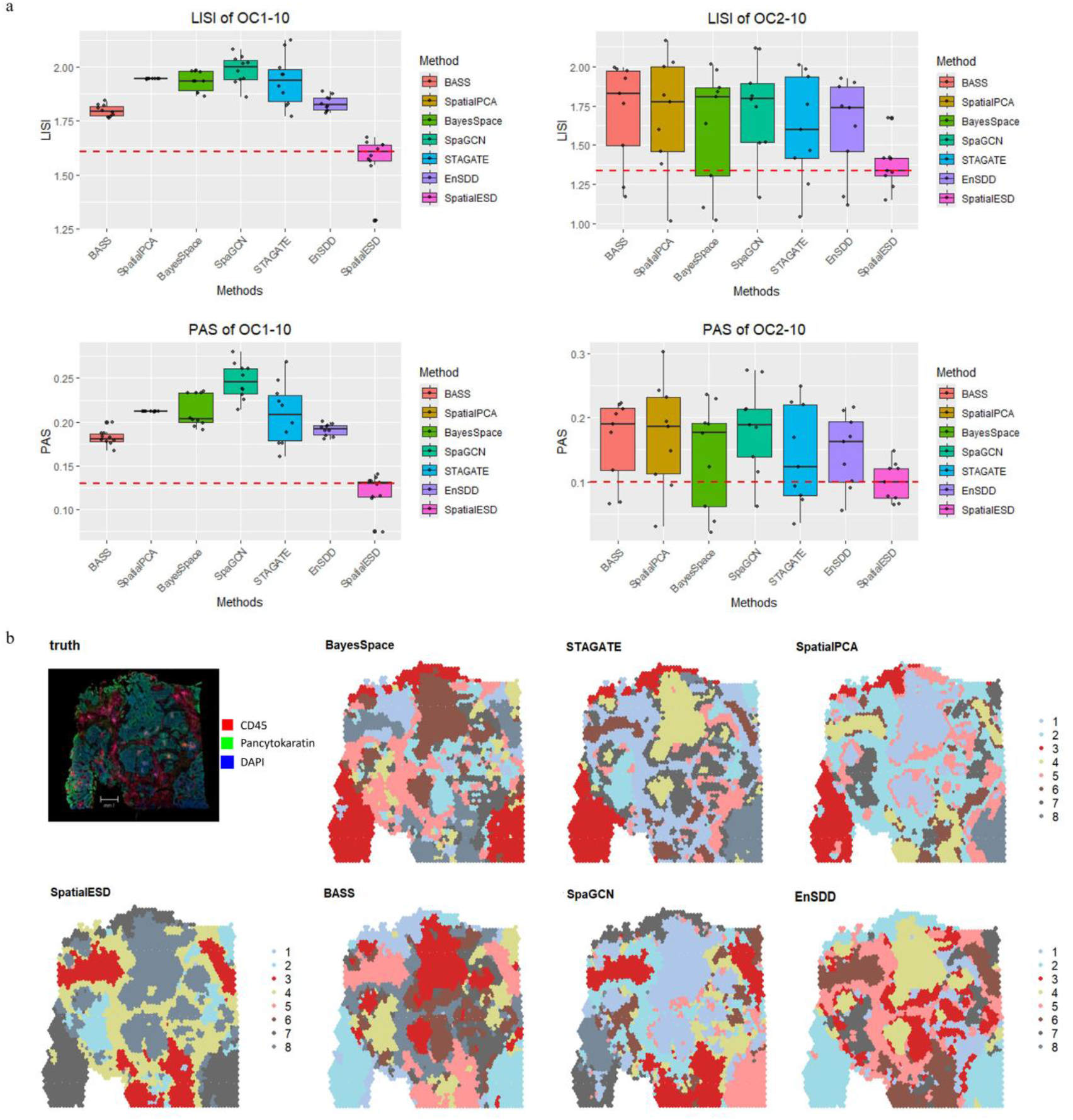
SpatialESD enhances tissue structure detection in ovarian cancer. (a) Boxplot results for the two evaluation metrics in the OC dataset under two different scenarios. (b) H&E-Stained Image and spatial domain visualizations of various methods.

Based on the spatial domains detected by SpatialESD, we did the Wilcoxon test (Figure 8a) and spatial autocorrelation analysis (Figure 8b), to identify domain-specific DEGs. Genes meeting the criteria were identified as DEGs, resulting in a total of 3,014 DEGs. Three CD45 region-specific DEGs were shown in Figure 8a. Differential gene analysis identified region-specific marker genes associated with distinct spatial domain types, such as *IGHG4, IGKC, MALAT1, MT-ATP, MT-ND2*, and *MTRNR2L*. Among these, *IGHG4, IGKC*, and *MALAT1* were predominantly enriched in CD45-labeled regions (representing leukocytes, mainly immune cells), while *MT-ATP* and *MT-ND2* showed enrichment in both CD45 and Pan-Cytokeratin regions. *MTRNR2L* was primarily enriched in Pan-Cytokeratin-labeled regions (representing epithelial cells). Previous studies have demonstrated that *IGHG4* is highly expressed in the tumor microenvironment (TME), potentially enhancing antitumor immune responses by regulating immune cell infiltration and activation, and is associated with improved patient prognosis ^[54]^. *IGHG4* influences immunotherapy efficacy by modulating B-cell and plasma cell-mediated antitumor immune responses ^[55]^. Overexpression of *MALAT1* accelerates tumor progression by promoting inflammatory responses, enhancing tumor cell proliferation and epithelial-mesenchymal transition (EMT), while suppressing apoptosis, making it a potential diagnostic and therapeutic target for epithelial ovarian cancer ^[56]^. Mutations in *MT-ATP* exhibit high mutation rates in various cancers, disrupting mitochondrial function and contributing to tumorigenesis and progression ^[57]^. *MT-ND2* mutations promote tumor recurrence by impairing mitochondrial function and driving the Warburg effect, conferring survival and drug resistance advantages ^[58]^. However, to our knowledge, no studies have reported the regulatory role of *MTRNR2L* in ovarian cancer, warranting further biological validation of these molecular markers. Figure 8b demonstrates that the top six genes with significantly differential expression in spatial domain category 6, compared to other domains, exhibited higher overall *G*_*i*_ values, indicating strong spatial autocorrelation.

**Figure 8.**
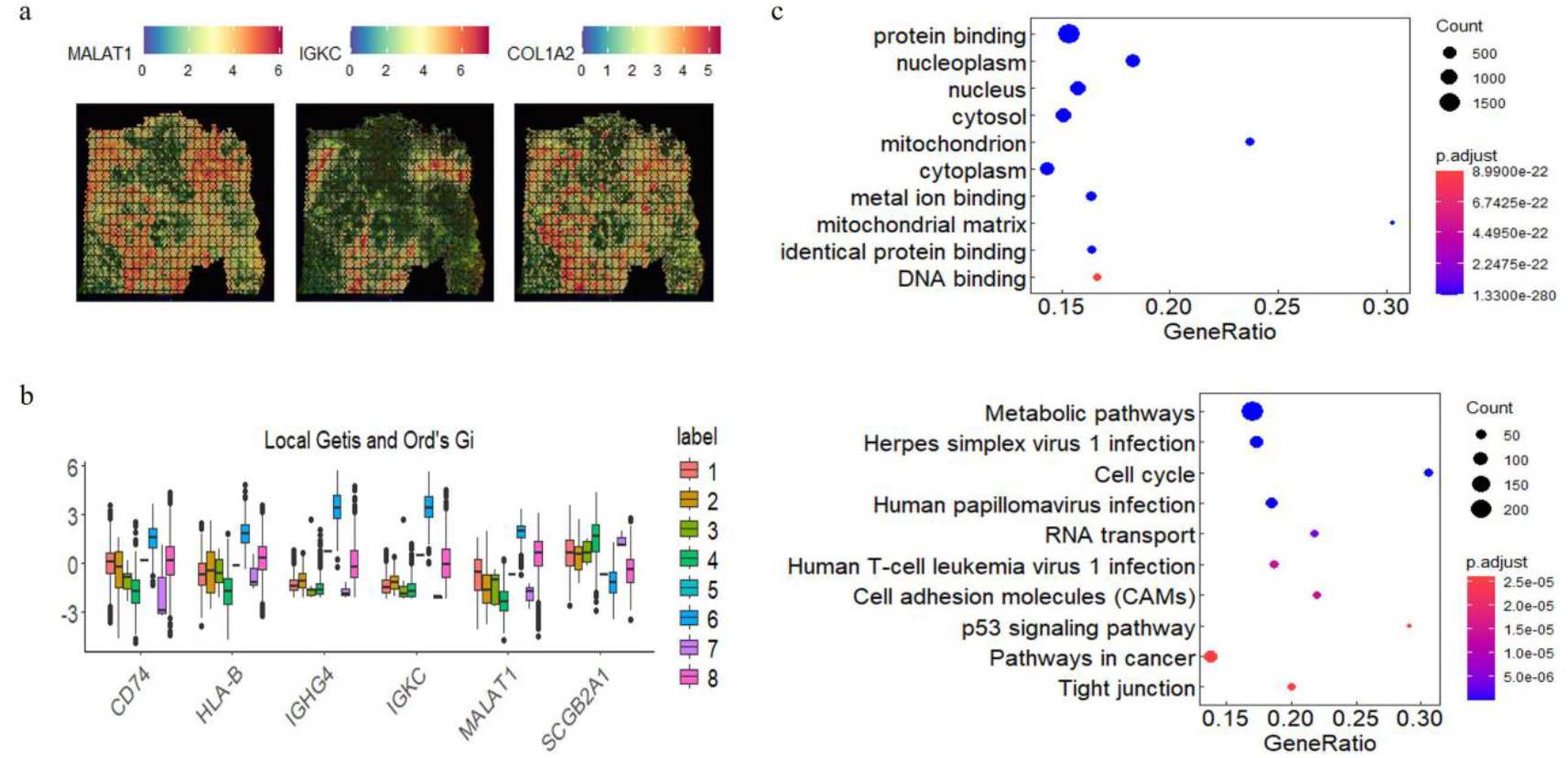
The downstream analysis corresponding to the OC dataset. (a) Visualization of differentially expressed genes selected using the Wilcoxon test. (b) Expression levels of differentially expressed genes in different spatial domains based on spatial autocorrelation analysis. (c) Enrichment analysis of differentially expressed genes (upper: GO enrichment; lower: KEGG pathway enrichment).

The 3,014 DEGs were subjected to functional enrichment analysis, including GO and KEGG analyses. The GO analysis (Figure 8c, upper panel) highlighted the top ten enriched biological processes and cellular components in OC, primarily involving critical intracellular functions and structures such as protein binding, metal ion binding, identical protein binding, and DNA binding (molecular functions); nucleoplasm, nucleus, cytosol, mitochondrion, cytoplasm, and mitochondrial matrix (cellular components), reflecting core roles in gene regulation, signal transduction, and metabolic homeostasis ^[59,60]^. KEGG analysis (Figure 8c, lower panel) revealed strong associations of DEGs with cancer-related pathways, including metabolic pathways, herpes simplex virus 1 infection, cell cycle, human papillomavirus infection, RNA transport, human T-cell leukemia virus 1 infection, cell adhesion molecules (CAMs), p53 signaling pathway, pathways in cancer, and tight junction. HSV-1, an oncolytic virus, selectively infects tumor cells and triggers immune responses in malignancies like ovarian cancer ^[61]^. Although weakly associated, human papillomavirus infection may contribute to ovarian carcinogenesis, particularly in regions with high prevalence, such as Asia ^[62]^. Alterations in the p53 signaling pathway vary across OC subtypes; for instance, p53 mutations are common in MAPK-negative high-grade serous carcinoma (HGSC) ^[63]^. The complete results of the downstream analysis are provided in Supplementary Note 3.

## 5 Discussion

In this study, we have proposed an enhanced ensemble learning method, SpatialESD, which aims to ensemble the results from different spatial clustering methods. By incorporating multi-scale spatial structural information, this method can capture spatial similarity in ST data with high precision. SpatialESD overcomes the limitations caused by the varying performance of different spatial domain detection methods on different datasets, addressing the challenge of selecting a specific method. In terms of clustering accuracy and stability, SpatialESD outperforms the five widely used advanced spatial domain detection methods, as well as the EnSDD ensemble method based on the same base clustering algorithms. Extensive validation on both simulated and real ST data demonstrates the practical effectiveness and advantages of SpatialESD.

Our method demonstrates significant innovation in two key aspects. First, we propose an ensemble approach specifically optimized for ST data, effectively improving the accuracy and stability of spatial domain detection to address the challenges in ST analysis. Second, by integrating multiple SDD methods and incorporating multi-scale spatial structural information, we ensure that our method delivers consistent and robust spatial domain detection results when applied to new datasets, enhancing its adaptability and reliability across different data environments.

In the simulation study, SpatialESD achieved higher ARI values and lower variability across datasets, demonstrating superior accuracy and stability in spatial domain detection. When applied to three annotated and two unannotated real datasets, SpatialESD substantially improved ARI values, with increases as high as 0.504 to 0.833. It also reduced the variability of clustering results, outperforming traditional base clustering methods in both accuracy and stability.

SpatialESD results showed strong concordance with manual annotations, validating its accuracy and robustness in spatial domain detection. Based on the spatial structures delineated by SpatialESD, we identified DEGs that displayed significant differential expression across specific domains. These DEGs reflect the distinct molecular features of the local tissue microenvironment and may regulate key processes such as tumor cell proliferation, migration, and immune evasion, underscoring their potential as clinical biomarkers. Functional enrichment analysis revealed that the identified DEGs are significantly associated with biological pathways central to tumor progression, including cell cycle regulation, extracellular matrix remodeling, and immune-related signaling. These findings further emphasize the critical roles of DEGs in tumor evolution.

Trajectory analysis was applied to reconstruct potential spatially resolved cell-state transitions, uncovering a dynamic progression from relatively quiescent to active cellular phenotypes within tumor regions. This provides new insights into the spatial mechanisms underlying tumor evolution. In addition, cell-cell communication analysis, integrated with reference datasets, enabled the transfer of cell-type annotations to the spatial transcriptomic data, achieving high-resolution detection of cellular subpopulations. This analysis revealed spatial interaction patterns among distinct cell types, facilitating the discovery of key ligand-receptor signaling pathways and a deeper understanding of cell-cell regulatory networks within the local microenvironment. The resulting communication networks offer valuable resources for identifying potential therapeutic targets, refining immunotherapeutic strategies, and discovering biomarkers for predicting disease progression and advancing precision medicine.

A limitation of the current SpatialESD method is its reliance solely on clustering outputs from base methods as input. Moreover, our study utilized only 10x Genomics spatial transcriptomics data, and the method has not yet been validated or adapted for other data types. Future work could focus on integrating a broader range of data sources, including both raw and processed data from diverse platforms, to further enhance performance and adaptability.

In conclusion, SpatialESD demonstrates clear advantages in capturing spatial similarity and accurately identifying spatial domains. Applied to IDC and OC datasets, the method successfully delineated spatial domains and associated biomarkers, providing a more precise and stable solution for ST analysis. By incorporating multi-scale spatial structural information, SpatialESD mitigates the performance variability observed in traditional methods across datasets, thereby improving spatial domain detection and offering valuable insights for tumor microenvironment research.

## Data availability

All datasets used in this paper are publicly available: (1) Human DLPFC data within the spatialLIBD (https://github.com/LieberInstitute/spatialLIBD); (2) Human breast cancer, ovarian cancer, invasive ductal carcinoma (https://www.10xgenomics.com/datasets); Human HER 2-positive breast cancer data. (https://github.com/almaan/her2st).

## Supplementary information

Supplementary material for this article is available online.

## Acknowledgments

The authors express their gratitude to the 10x genomic Program for maintaining crucial public databases and services.

## Funding

National Natural Science Foundation of China [82473739 to H.C., 82273742 to H.Y.]; Applied Basic Research Project of Shanxi Province [202303021211130]; Shanxi Province Research Funding Project for Returned Overseas Scholars [2024-081]; Shanxi Province Higher Education “Billion Project” Science and Technology Guidance Project, Open Project Fund from Key Laboratory of Coal Environmental Pathogenicity and Prevention (Shanxi Medical University), Ministry of Education, China [MEKLCEPP/SXMU-202415] and a fund from Michigan State University (to Y.C.).

## Author Contributions

H.C. performed the study; H.C. and Y.C. conceived the idea; G.L., J.X., R.C., and T.W. assisted with the real data analysis; X.Y., R.F., P.Z., Y.L., H.Y. and Y.Z. contributed to the interpretation of the results; H.C. and Y.C. wrote the manuscript with input from all other authors. All authors reviewed and approved the final manuscript.

## Conflict of interest statement

The authors declare no competing interests.

## Notes

### Competing Interest Statement

The authors have declared no competing interest.

