## Supplementary File for "SpatialESD: spatial ensemble domain detection in spatial transcriptomics"

### Supplementary Information for "SpatialESD: spatial ensemble domain detection in spatial transcriptomics"

#### 1. Supplementary Tables:

**Table S1** Summary of Key Spatial Domain Detection Methods

| Methods | Keyword descriptions | Histology | Website | Language |
|---|---|---|---|---|
| BayesSpace | Bayesian model with a Markov random field | × | https://www.bioconductor.org/packages/release/bioc/html/BayesSpace | R |
| BASS | multi-scale and multi-sample analysis | × | https://github.com/zhengli09/BASS | R |
| SpaGCN | graph convolutional network-based model | √ | https://github.com/jianhuupenn/SpaGCN | Python |
| STAGATE | graph attention auto-encoder framework | √ | https://github.com/ttgump/spaVAE/tree/main/src/spaVAE | Python |
| SpatialPCA | Spatially aware dimension reduction | × | https://github.com/shangll123/SpatialPCA. | R |

**Table S2** ARI Results of Different SDD Methods for simulated data

| | BayesSpace | BASS | SpaGCN | STAGATE | SpatialPCA | EnSDD | SpatialESD |
|---|---|---|---|---|---|---|---|
| 01 | 0.369 | 0.227 | 0.196 | 0.433 | 0.491 | 0.402 | 0.475 |
| 02 | 0.245 | 0.213 | 0.362 | 0.462 | 0.366 | 0.314 | **0.500** |
| 03 | 0.390 | 0.322 | 0.186 | 0.478 | 0.366 | 0.329 | **0.503** |
| 04 | 0.371 | 0.288 | 0.213 | 0.362 | 0.363 | 0.321 | **0.479** |
| 05 | 0.223 | 0.203 | 0.347 | 0.444 | 0.379 | 0.357 | 0.390 |
| 06 | 0.219 | 0.300 | 0.314 | 0.444 | 0.662 | 0.393 | 0.550 |
| 07 | 0.388 | 0.233 | 0.363 | 0.220 | 0.658 | 0.319 | 0.542 |
| 08 | 0.385 | 0.210 | 0.216 | 0.433 | 0.534 | 0.267 | 0.479 |
| 09 | 0.383 | 0.216 | 0.208 | 0.456 | 0.669 | 0.387 | 0.523 |
| 10 | 0.384 | 0.210 | 0.290 | 0.360 | 0.509 | 0.340 | **0.511** |
| 11 | 0.369 | 0.210 | 0.222 | 0.369 | 0.344 | 0.404 | **0.483** |

| | | | | | | | |
|---|---|---|---|---|---|---|---|
| 12 | 0.194 | 0.231 | 0.363 | 0.425 | 0.495 | 0.323 | 0.356 |
| 13 | 0.217 | 0.217 | 0.365 | 0.202 | 0.376 | 0.308 | **0.534** |
| 14 | 0.238 | 0.227 | 0.188 | 0.314 | 0.395 | 0.285 | **0.523** |
| 15 | 0.385 | 0.205 | 0.324 | 0.285 | 0.394 | 0.323 | **0.695** |
| 16 | 0.394 | 0.228 | 0.226 | 0.430 | 0.516 | 0.261 | 0.367 |
| 17 | 0.368 | 0.211 | 0.351 | 0.439 | 0.382 | 0.288 | **0.523** |
| 18 | 0.245 | 0.347 | 0.192 | 0.362 | 0.366 | 0.323 | **0.722** |
| 19 | 0.387 | 0.228 | 0.369 | 0.365 | 0.391 | 0.410 | **0.540** |
| 20 | 0.377 | 0.222 | 0.353 | 0.442 | 0.677 | 0.414 | 0.535 |

**Table S3** ARI Results of Different SDD Methods for DLPFC

| | BayesSpace | BASS | SpaGCN | STAGATE | SpatialPCA | EnSDD | SpatialESD |
|---|---|---|---|---|---|---|---|
| 151507 | 0.468 | 0.447 | 0.471 | 0.538 | 0.539 | 0.538 | 0.562 |
| 151508 | 0.437 | 0.406 | 0.395 | 0.456 | 0.453 | 0.508 | 0.317 |
| 151509 | 0.381 | 0.431 | 0.459 | 0.325 | 0.554 | 0.463 | 0.435 |
| 151510 | 0.374 | 0.420 | 0.430 | 0.474 | 0.419 | 0.474 | 0.533 |
| 151669 | 0.468 | 0.382 | 0.282 | 0.451 | 0.374 | 0.469 | 0.542 |
| 151670 | 0.428 | 0.378 | 0.369 | 0.439 | 0.519 | 0.385 | 0.69 |
| 151671 | 0.730 | 0.556 | 0.504 | 0.590 | 0.595 | 0.571 | 0.833 |
| 151672 | 0.426 | 0.556 | 0.557 | 0.589 | 0.527 | 0.583 | 0.601 |
| 151673 | 0.546 | 0.579 | 0.515 | 0.594 | 0.571 | 0.591 | 0.658 |
| 151674 | 0.291 | 0.637 | 0.385 | 0.557 | 0.547 | 0.626 | 0.596 |
| 151675 | 0.525 | 0.606 | 0.352 | 0.580 | 0.544 | 0.640 | 0.610 |
| 151676 | 0.364 | 0.607 | 0.328 | 0.426 | 0.629 | 0.488 | 0.428 |

**Table S4** ARI Results of Different SDD Methods for Breast cancer

| | BayesSpace | SpaGCN | STAGATE | SpatialPCA | EnSDD | SpatialESD |
|---|---|---|---|---|---|---|
| Run1 | 0.500 | 0.542 | 0.456 | 0.319 | 0.566 | 0.603 |
| Run2 | 0.446 | 0.540 | 0.459 | 0.319 | 0.566 | 0.607 |
| Run3 | 0.505 | 0.512 | 0.556 | 0.319 | 0.578 | 0.591 |
| Run4 | 0.493 | 0.482 | 0.531 | 0.319 | 0.591 | 0.499 |
| Run5 | 0.455 | 0.500 | 0.553 | 0.319 | 0.684 | 0.582 |
| Run6 | 0.480 | 0.503 | 0.510 | 0.319 | 0.579 | 0.587 |
| Run7 | 0.479 | 0.508 | 0.554 | 0.319 | 0.474 | 0.569 |
| Run8 | 0.501 | 0.494 | 0.485 | 0.319 | 0.511 | 0.587 |
| Run9 | 0.494 | 0.509 | 0.513 | 0.319 | 0.515 | 0.599 |
| Run10 | 0.506 | 0.475 | 0.468 | 0.319 | 0.549 | 0.606 |

**Table S5** ARI Results of Different SDD Methods for HER2

| | BayesSpace | BASS | SpaGCN | STAGATE | SpatialPCA | EnSDD | SpatialESD |
|---|---|---|---|---|---|---|---|
| A1 | 0.271 | 0.412 | 0.107 | 0.313 | 0.142 | 0.218 | 0.226 |
| B1 | 0.199 | 0.342 | 0.221 | 0.221 | 0.276 | 0.314 | 0.257 |
| C1 | 0.309 | 0.507 | 0.099 | -0.106 | -0.150 | 0.297 | **0.514** |
| D1 | 0.203 | 0.193 | 0.182 | 0.130 | 0.103 | 0.207 | **0.203** |
| E1 | 0.011 | 0.119 | 0.081 | 0.246 | -0.077 | 0.126 | 0.011 |
| F1 | 0.115 | 0.079 | 0.078 | 0.022 | 0.147 | 0.115 | **0.300** |
| G2 | 0.098 | 0.220 | 0.165 | 0.147 | 0.182 | 0.201 | 0.160 |
| H1 | 0.292 | 0.296 | 0.267 | 0.293 | 0.308 | 0.306 | 0.259 |

**Table S6** LISI Results of Different SDD Methods for IDC (Number of Clusters from 2 to 10)

| Cluster | BayesSpace | BASS | SpaGCN | STAGATE | SpatialPCA | EnSDD | SpatialESD |
|---|---|---|---|---|---|---|---|
| 2 | 1.000 | 1.166 | 1.000 | 1.000 | 1.000 | 1.000 | 1.000 |
| 3 | 1.000 | 1.063 | 1.059 | 1.000 | 1.000 | 1.000 | 1.000 |
| 4 | 1.065 | 1.160 | 1.115 | 1.008 | 1.002 | 1.076 | 1.000 |
| 5 | 1.158 | 1.199 | 1.129 | 1.032 | 1.025 | 1.088 | 1.000 |
| 6 | 1.098 | 1.281 | 1.181 | 1.168 | 1.082 | 1.076 | 1.002 |
| 7 | 1.204 | 1.324 | 1.183 | 1.183 | 1.298 | 1.208 | 1.062 |
| 8 | 1.200 | 1.400 | 1.388 | 1.260 | 1.492 | 1.298 | 1.041 |
| 9 | 1.367 | 1.394 | 1.418 | 1.274 | 1.510 | 1.298 | 1.099 |
| 10 | 1.447 | 1.432 | 1.386 | 1.317 | 1.515 | 1.386 | 1.170 |

**Table S7** PAS Results of Different SDD Methods for IDC (Number of Clusters from 2 to 10)

| Cluster | BayesSpace | BASS | SpaGCN | STAGATE | SpatialPCA | EnSDD | SpatialESD |
|---|---|---|---|---|---|---|---|
| 2 | 0.002 | 0.048 | 0.016 | 0.003 | 0.006 | 0.003 | 0.003 |
| 3 | 0.008 | 0.033 | 0.029 | 0.005 | 0.009 | 0.007 | 0.006 |
| 4 | 0.027 | 0.066 | 0.049 | 0.011 | 0.011 | 0.030 | 0.008 |
| 5 | 0.040 | 0.067 | 0.050 | 0.015 | 0.017 | 0.045 | 0.016 |
| 6 | 0.041 | 0.088 | 0.071 | 0.035 | 0.028 | 0.029 | 0.017 |
| 7 | 0.055 | 0.094 | 0.074 | 0.048 | 0.058 | 0.094 | 0.037 |
| 8 | 0.058 | 0.099 | 0.110 | 0.065 | 0.112 | 0.094 | 0.032 |
| 9 | 0.077 | 0.097 | 0.119 | 0.063 | 0.113 | 0.08 | 0.041 |
| 10 | 0.111 | 0.110 | 0.112 | 0.072 | 0.115 | 0.092 | 0.073 |

**Table S8** LISI Results of Different SDD Methods for IDC under 10 Random Seeds

| Seed | BayesSpace | BASS | SpaGCN | STAGATE | SpatialPCA | EnSDD | SpatialESD |
|---|---|---|---|---|---|---|---|
| 1 | 1.453 | 1.432 | 1.479 | 1.307 | 1.498 | 1.316 | 1.133 |
| 2 | 1.416 | 1.437 | 1.517 | 1.325 | 1.498 | 1.286 | 1.169 |
| 3 | 1.425 | 1.37 | 1.443 | 1.386 | 1.498 | 1.293 | 1.204 |
| 4 | 1.394 | 1.433 | 1.303 | 1.290 | 1.498 | 1.298 | 1.147 |
| 5 | 1.419 | 1.373 | 1.412 | 1.278 | 1.498 | 1.269 | 1.162 |
| 6 | 1.446 | 1.442 | 1.459 | 1.311 | 1.498 | 1.302 | 1.189 |
| 7 | 1.383 | 1.441 | 1.444 | 1.274 | 1.498 | 1.313 | 1.042 |
| 8 | 1.383 | 1.427 | 1.465 | 1.277 | 1.498 | 1.282 | 1.179 |
| 9 | 1.392 | 1.434 | 1.487 | 1.437 | 1.498 | 1.351 | 1.306 |
| 10 | 1.419 | 1.353 | 1.334 | 1.324 | 1.498 | 1.29 | 1.184 |

**Table S9** PAS Results of Different SDD Methods for IDC under 10 Random Seeds

| Seed | BayesSpace | BASS | SpaGCN | STAGATE | SpatialPCA | EnSDD | SpatialESD |
|---|---|---|---|---|---|---|---|
| 1 | 0.109 | 0.11 | 0.119 | 0.074 | 0.094 | 0.085 | 0.058 |
| 2 | 0.092 | 0.111 | 0.125 | 0.078 | 0.094 | 0.076 | 0.077 |
| 3 | 0.084 | 0.095 | 0.117 | 0.082 | 0.094 | 0.074 | 0.056 |
| 4 | 0.100 | 0.112 | 0.099 | 0.064 | 0.094 | 0.094 | 0.066 |
| 5 | 0.092 | 0.100 | 0.115 | 0.066 | 0.094 | 0.072 | 0.080 |
| 6 | 0.099 | 0.114 | 0.102 | 0.073 | 0.094 | 0.084 | 0.050 |
| 7 | 0.08 | 0.110 | 0.135 | 0.067 | 0.094 | 0.098 | 0.029 |
| 8 | 0.081 | 0.112 | 0.123 | 0.076 | 0.094 | 0.082 | 0.054 |
| 9 | 0.099 | 0.113 | 0.135 | 0.104 | 0.094 | 0.117 | 0.086 |
| 10 | 0.092 | 0.107 | 0.078 | 0.072 | 0.094 | 0.079 | 0.048 |

**Table S10** LISI Results of Different SDD Methods for OC (Number of Clusters from 2 to 10)

| Cluster | BayesSpace | BASS | SpaGCN | STAGATE | SpatialPCA | EnSDD | SpatialESD |
|---|---|---|---|---|---|---|---|
| 2 | 1.021 | 1.172 | 1.168 | 1.043 | 1.020 | 1.119 | 1.149 |
| 3 | 1.101 | 1.232 | 1.512 | 1.251 | 1.379 | 1.175 | 1.237 |
| 4 | 1.307 | 1.498 | 1.516 | 1.415 | 1.460 | 1.458 | 1.326 |
| 5 | 1.637 | 1.763 | 1.793 | 1.463 | 1.601 | 1.619 | 1.336 |
| 6 | 1.804 | 1.829 | 1.745 | 1.600 | 1.774 | 1.735 | 1.307 |
| 7 | 1.836 | 1.921 | 1.892 | 1.935 | 1.819 | 1.748 | 1.673 |
| 8 | 1.863 | 1.993 | 1.814 | 1.982 | 1.997 | 1.895 | 1.419 |
| 9 | 1.979 | 1.980 | 2.108 | 1.76 | 2.027 | 1.869 | 1.414 |
| 10 | 2.011 | 1.969 | 2.116 | 2.01 | 2.163 | 1.926 | 1.421 |

**Table S11** PAS Results of Different SDD Methods for OC (Number of Clusters from 2 to 10)

| Cluster | BayesSpace | BASS | SpaGCN | STAGATE | SpatialPCA | EnSDD | SpatialESD |
|---|---|---|---|---|---|---|---|
| 2 | 0.022 | 0.068 | 0.061 | 0.035 | 0.030 | 0.055 | 0.064 |
| 3 | 0.038 | 0.065 | 0.115 | 0.072 | 0.094 | 0.090 | 0.099 |
| 4 | 0.062 | 0.118 | 0.139 | 0.093 | 0.112 | 0.100 | 0.075 |
| 5 | 0.123 | 0.177 | 0.185 | 0.079 | 0.148 | 0.127 | 0.077 |
| 6 | 0.176 | 0.190 | 0.188 | 0.123 | 0.186 | 0.162 | 0.066 |
| 7 | 0.190 | 0.206 | 0.212 | 0.220 | 0.193 | 0.170 | 0.148 |
| 8 | 0.191 | 0.223 | 0.213 | 0.224 | 0.231 | 0.210 | 0.121 |
| 9 | 0.229 | 0.220 | 0.271 | 0.169 | 0.242 | 0.193 | 0.121 |
| 10 | 0.236 | 0.214 | 0.273 | 0.249 | 0.302 | 0.216 | 0.127 |

**Table S12** LISI Results of Different SDD Methods for OC under 10 Random Seeds

| Seed | BayesSpace | BASS | SpaGCN | STAGATE | SpatialPCA | EnSDD | SpatialESD |
|---|---|---|---|---|---|---|---|
| 1 | 1.982 | 1.820 | 1.862 | 1.996 | 1.945 | 1.856 | 1.673 |
| 2 | 1.879 | 1.766 | 2.036 | 1.773 | 1.945 | 1.794 | 1.590 |
| 3 | 1.977 | 1.780 | 2.018 | 1.830 | 1.945 | 1.799 | 1.651 |
| 4 | 1.863 | 1.763 | 1.942 | 1.882 | 1.945 | 1.817 | 1.573 |
| 5 | 1.935 | 1.799 | 2.047 | 1.965 | 1.945 | 1.877 | 1.543 |
| 6 | 1.982 | 1.843 | 1.931 | 2.126 | 1.945 | 1.889 | 1.637 |
| 7 | 1.937 | 1.824 | 2.020 | 1.910 | 1.945 | 1.786 | 1.621 |
| 8 | 1.935 | 1.809 | 1.984 | 1.967 | 1.945 | 1.831 | 1.638 |
| 9 | 1.984 | 1.773 | 1.945 | 2.101 | 1.945 | 1.855 | 1.289 |
| 10 | 1.879 | 1.79 | 2.083 | 1.821 | 1.945 | 1.813 | 1.563 |

**Table S13** PAS Results of Different SDD Methods for OC under 10 Random Seeds

| Seed | BayesSpace | BASS | SpaGCN | STAGATE | SpatialPCA | EnSDD | SpatialESD |
|---|---|---|---|---|---|---|---|
| 1 | 0.233 | 0.182 | 0.214 | 0.219 | 0.212 | 0.195 | 0.140 |
| 2 | 0.199 | 0.178 | 0.253 | 0.160 | 0.212 | 0.182 | 0.129 |
| 3 | 0.233 | 0.184 | 0.280 | 0.176 | 0.212 | 0.180 | 0.137 |
| 4 | 0.191 | 0.176 | 0.237 | 0.198 | 0.212 | 0.190 | 0.113 |
| 5 | 0.202 | 0.178 | 0.266 | 0.232 | 0.212 | 0.198 | 0.115 |
| 6 | 0.235 | 0.200 | 0.231 | 0.248 | 0.212 | 0.200 | 0.132 |
| 7 | 0.204 | 0.180 | 0.261 | 0.188 | 0.212 | 0.184 | 0.132 |
| 8 | 0.203 | 0.187 | 0.238 | 0.224 | 0.212 | 0.196 | 0.131 |
| 9 | 0.233 | 0.167 | 0.226 | 0.269 | 0.212 | 0.194 | 0.075 |
| 10 | 0.195 | 0.188 | 0.261 | 0.175 | 0.212 | 0.190 | 0.115 |

#### 2. Supplementary Figures:

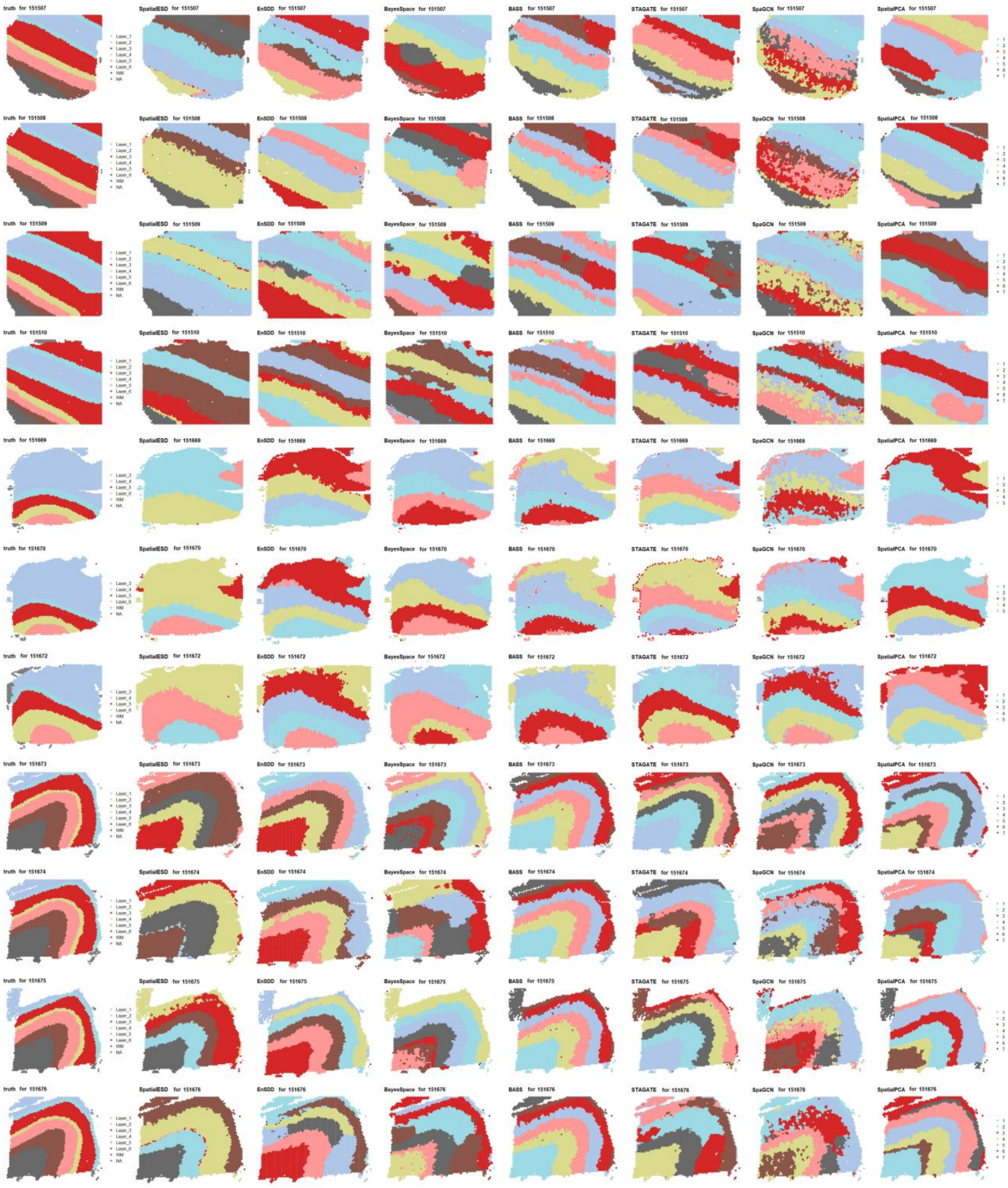

**Figure S1.** Spatial domain visualization results for different methods on 12 slices of the Human Dorsolateral Prefrontal Cortex Dataset.

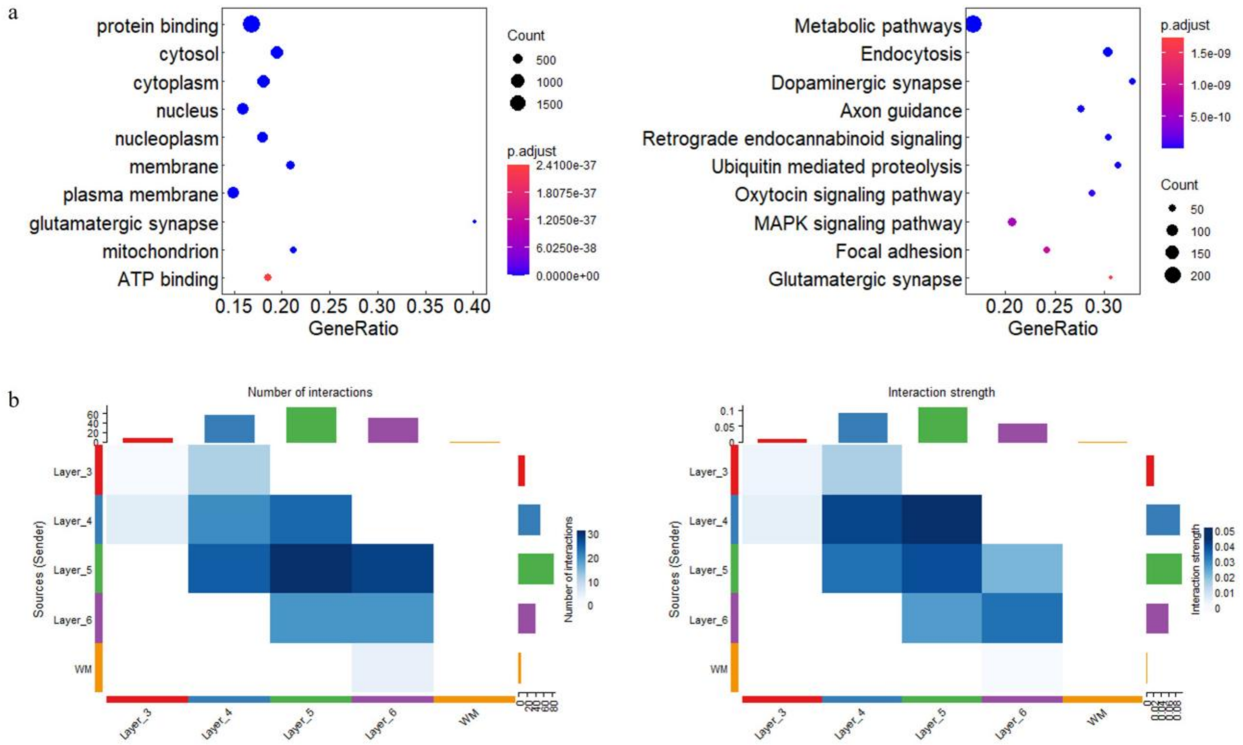


**Figure S2.** (a) Enrichment Analysis of Slice 151671 in DLPFC: Left: GO Analysis, Right: KEGG Analysis. (b) Number of Interactions and Interaction Strength.

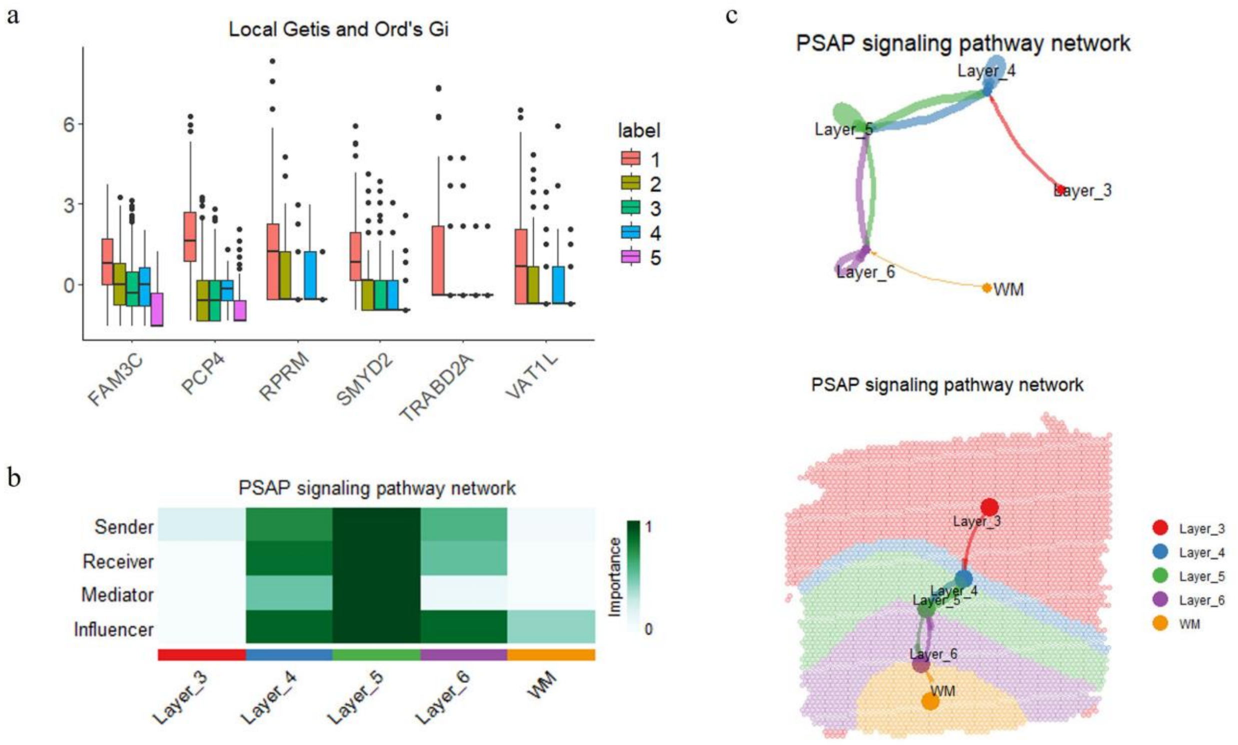


**Figure S3.** (a) Expression levels of differentially expressed genes in different spatial domains based on spatial autocorrelation analysis for 151671. (b) Heatmap of Centrality Scores in PSAP Signaling Pathway Network for 151671. (c) Circle plot and spatial map with signaling overlay of the PSAP Signaling Pathway Network for 151671.

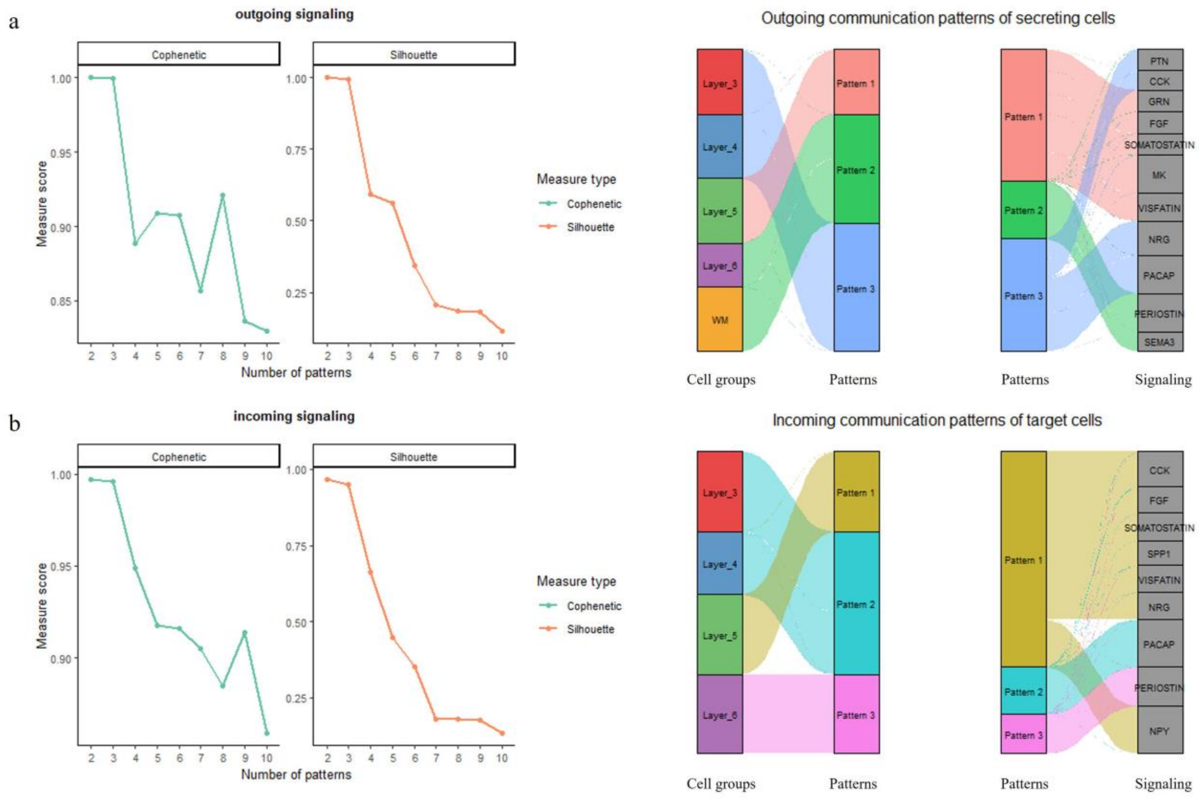


**Figure S4.** (a) Outgoing Communication Patterns in Slice 151671. Left: Determination of the number of inferred outgoing communication patterns; Right: River plot showing outgoing communication patterns of secreting cells. (b) Incoming Communication Patterns in Slice 151671. Left: Determination of the number of inferred incoming communication patterns; Right: River plot showing incoming communication patterns of secreting cells.

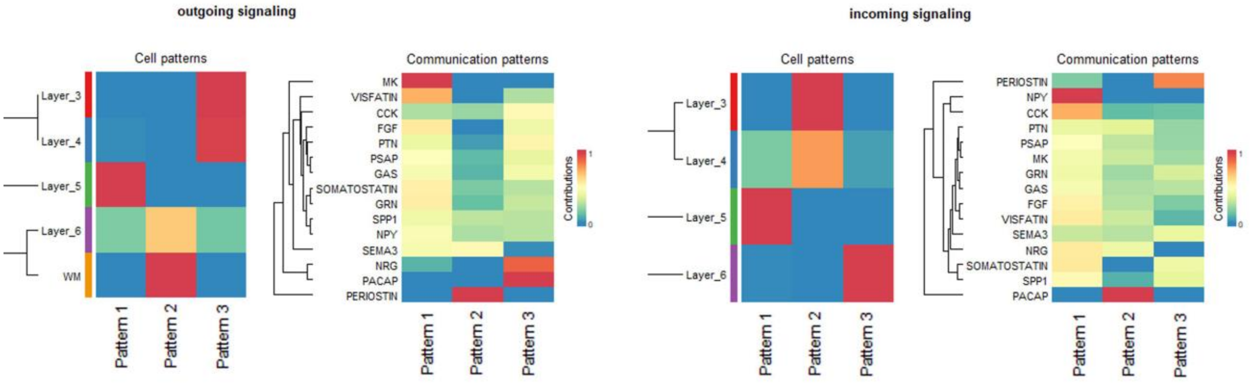


**Figure S5.** Cell and Communication Pattern Contributions in Outgoing and Incoming Signaling for 151671.

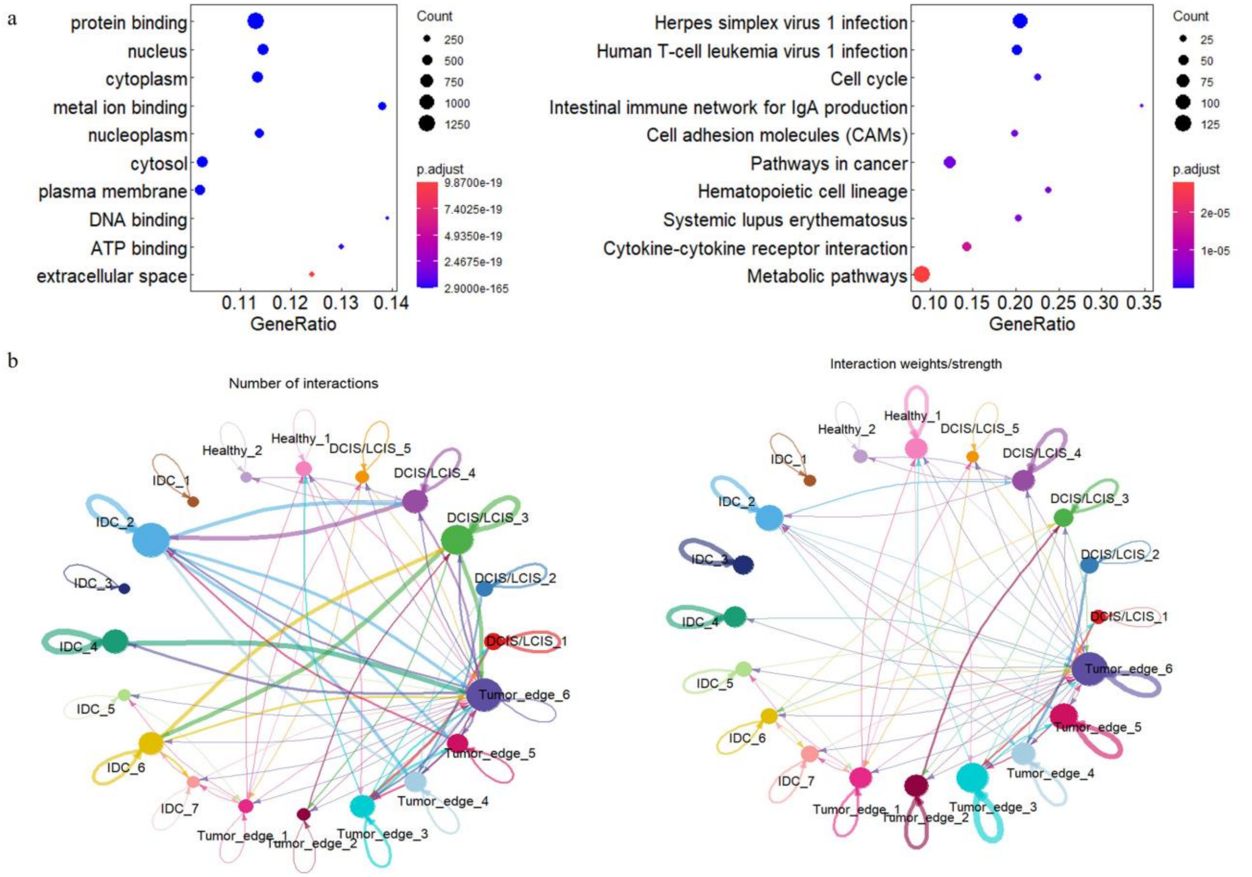


**Figure S6.** (a) Enrichment Analysis of breast cancer: Left: GO Analysis; Right: KEGG Analysis. (b) Number of Interactions and Interaction Strength of breast cancer.

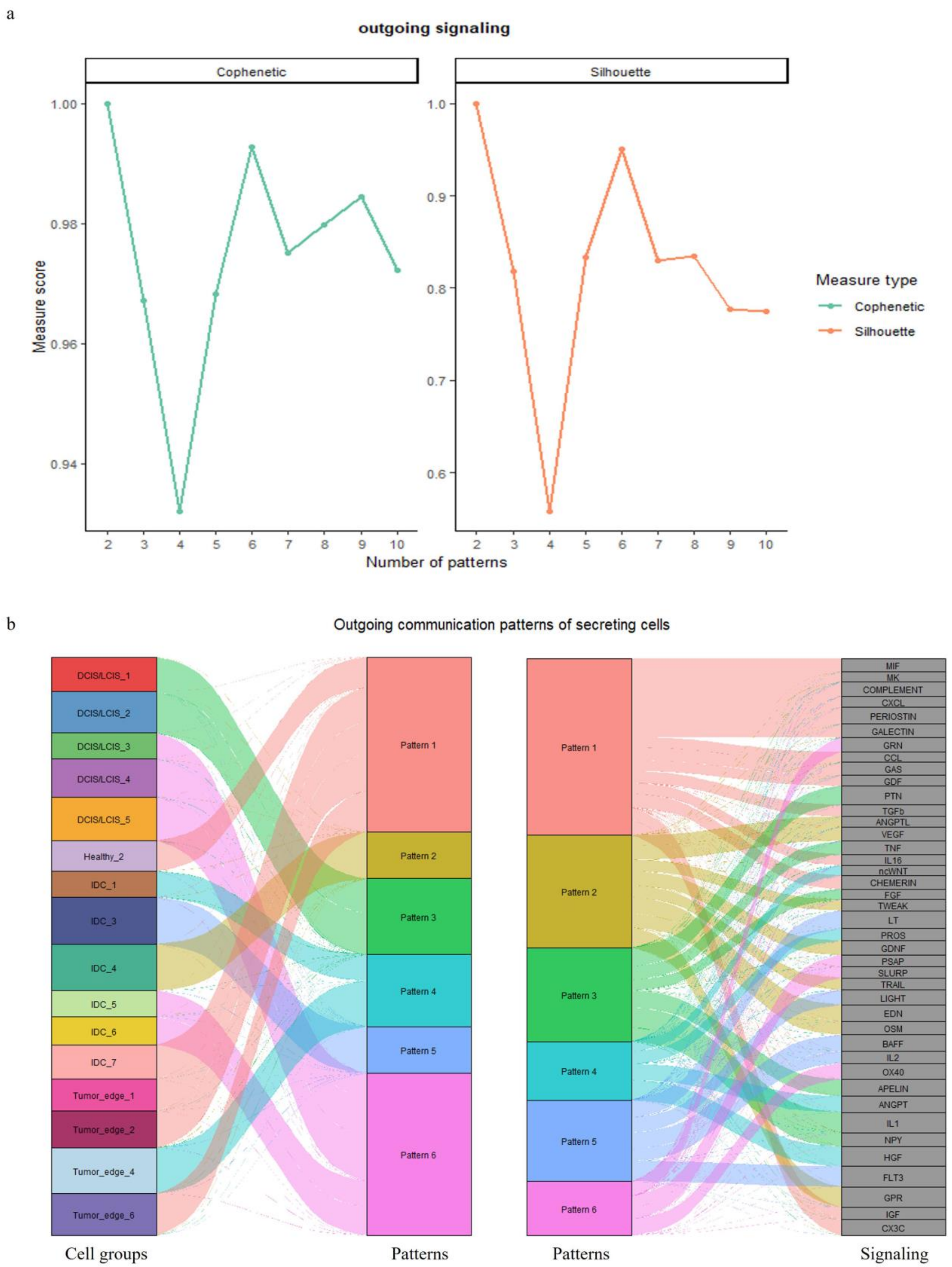


**Figure S7.** (a) Determination of the number of inferred outgoing communication patterns for breast cancer. (b) River plot showing outgoing communication patterns of secreting cells.

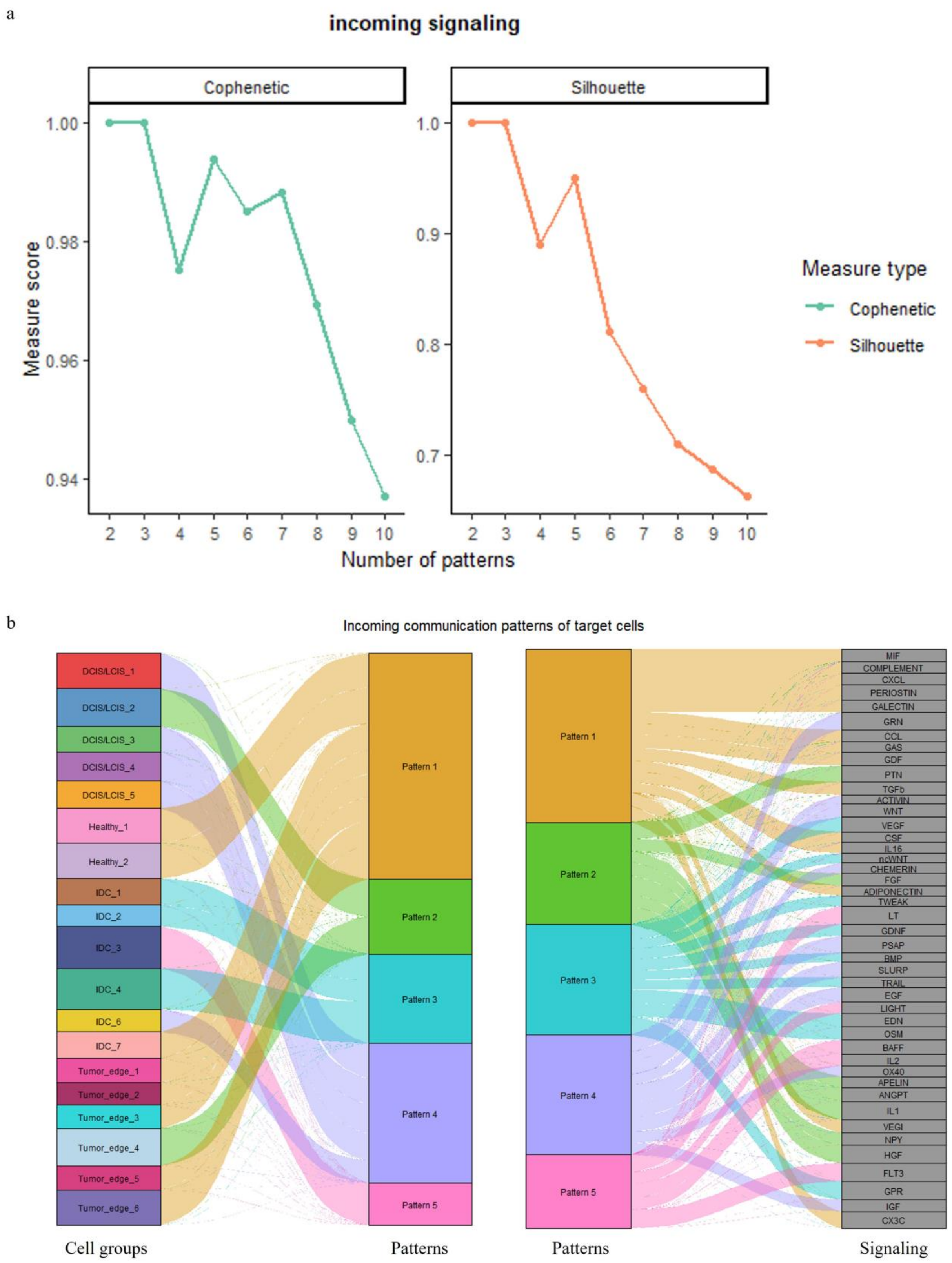


**Figure S8.** (a) Determination of the number of inferred incoming communication patterns for breast cancer. (b) River plot showing incoming communication patterns of secreting cells.

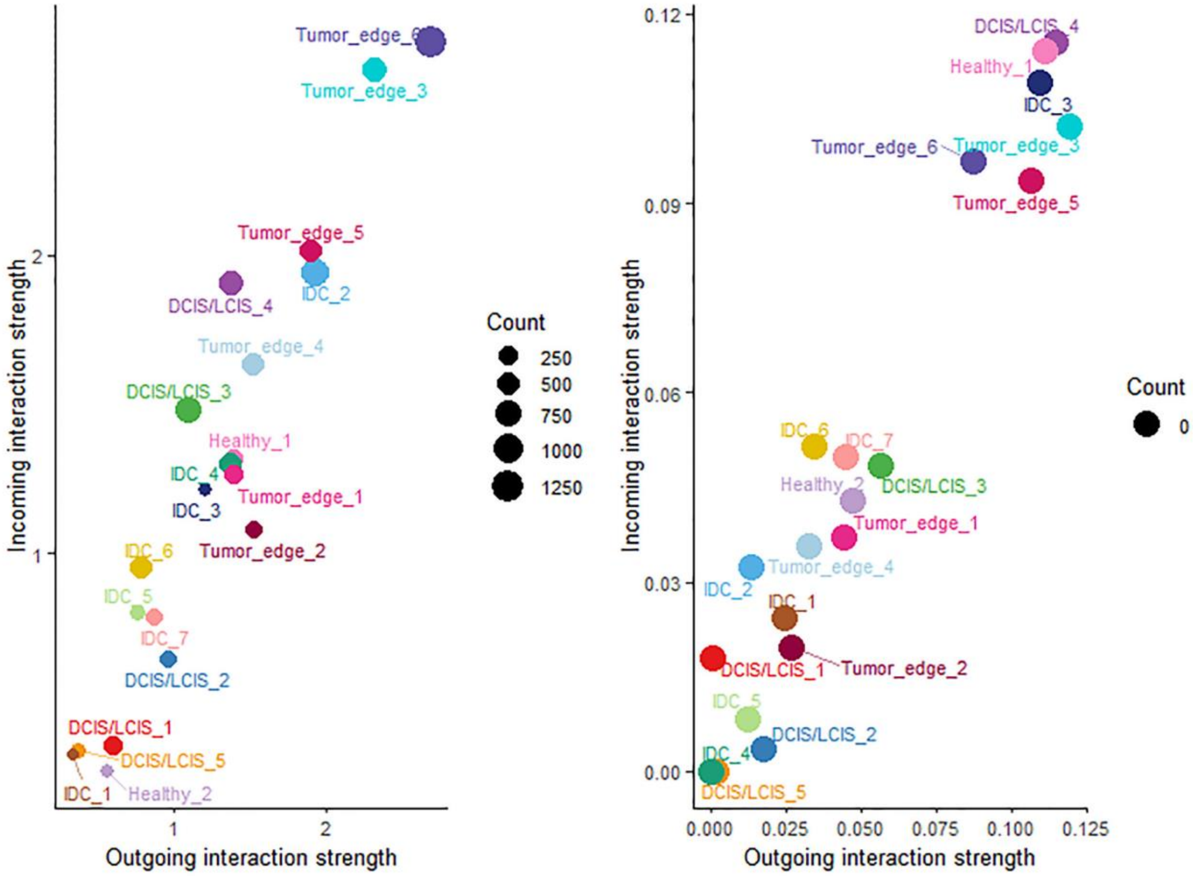


Figure S9. Global and CXCL Specific Signaling Role Analysis in Cell-Cell Communication for breast cancer.

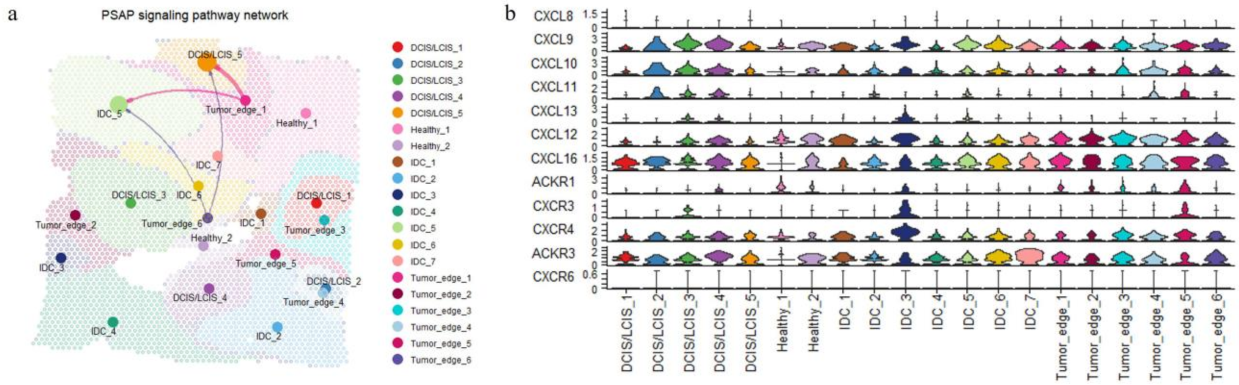


**Figure S10.** (a) Spatial map with signaling overlay of the PSAP Signaling Pathway Network for breast cancer. (b) Gene Expression in CXCL Signaling Pathway for breast cancer.

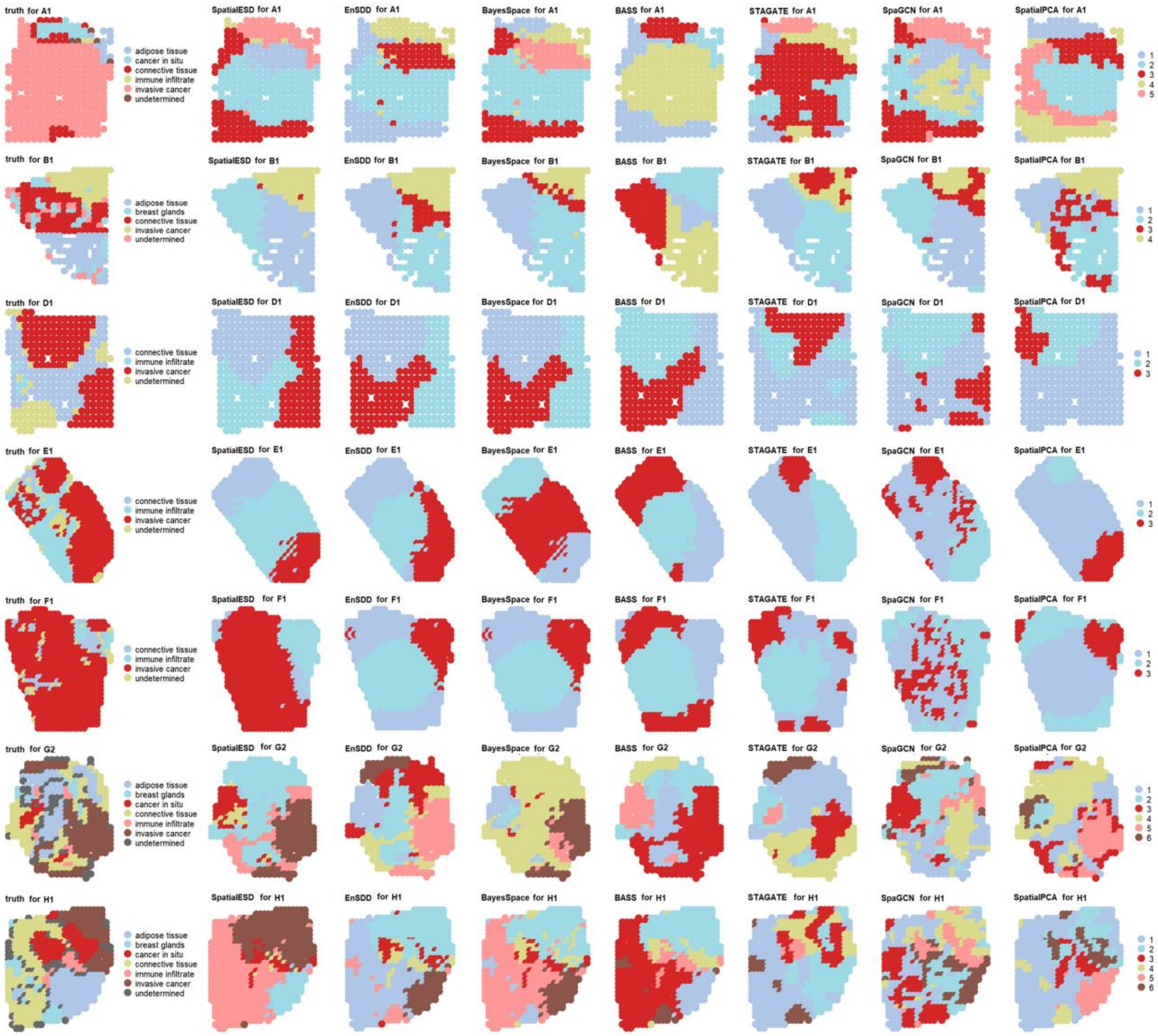


**Figure S11.** Spatial domain visualization results for different methods on 8 slices of the HER2.

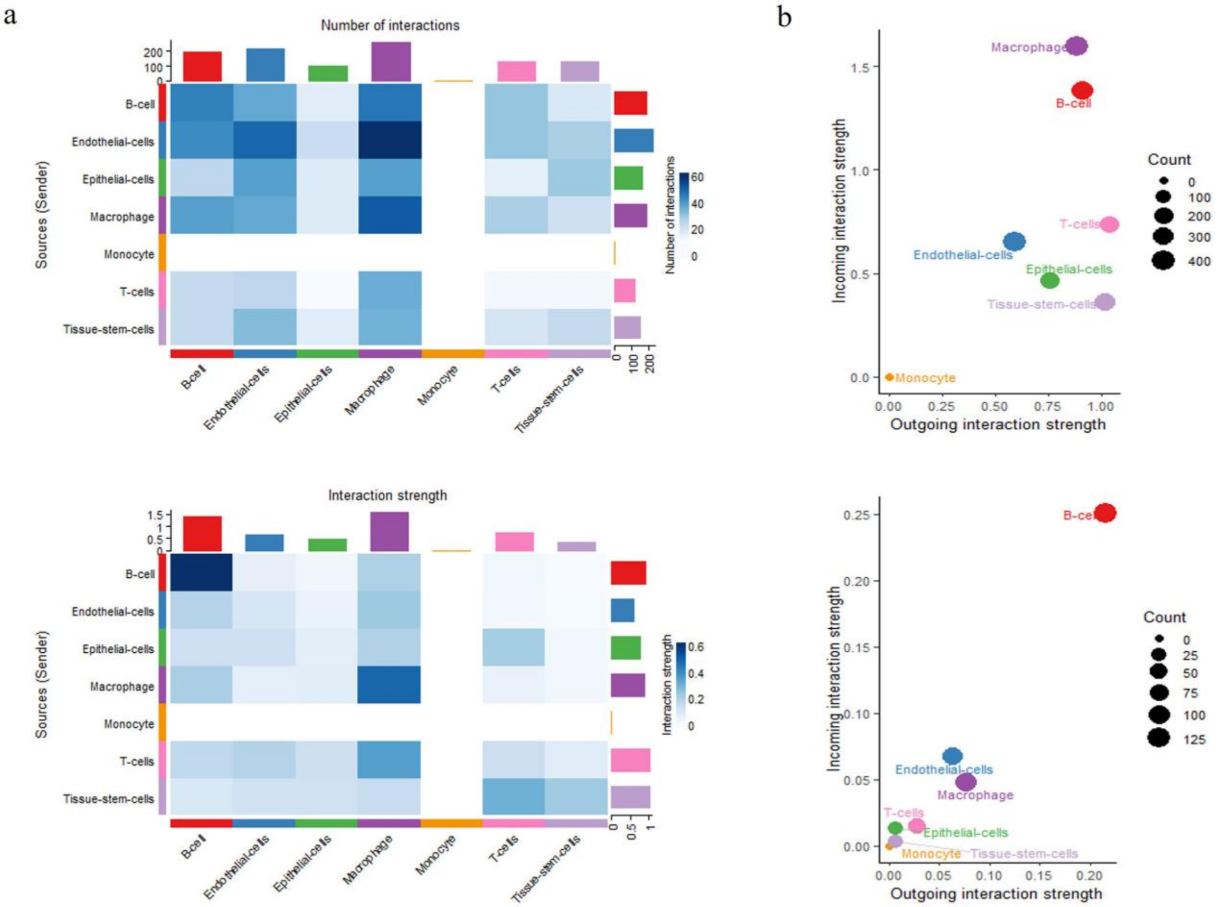


**Figure S12.** (a) Number of Interactions and Interaction Strength for IDC. (b) Global and CXCL/CCL- Specific Signaling Role Analysis in Cell-Cell Communication for IDC.

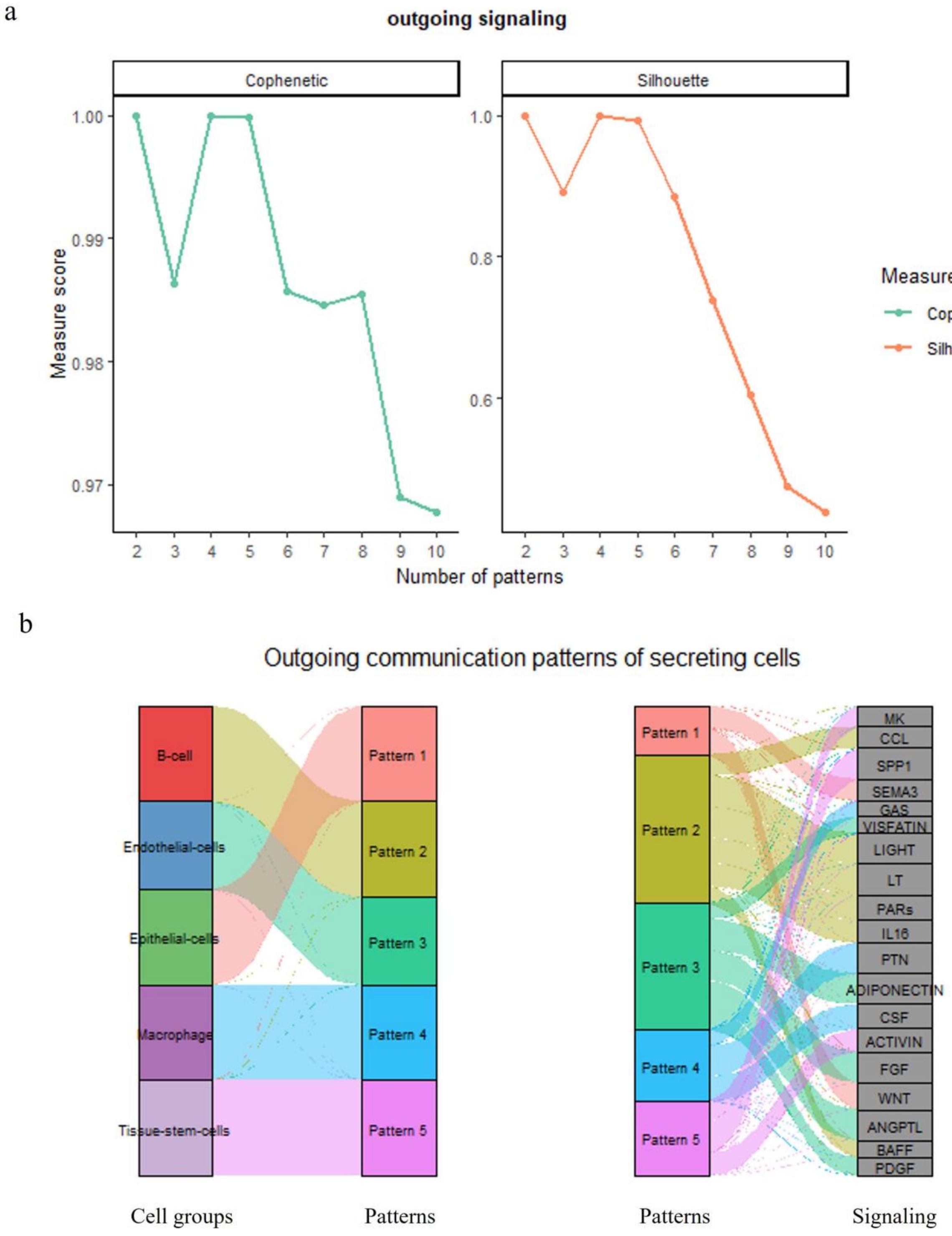


**Figure S13.** (a) Determination of the number of inferred outgoing communication patterns for IDC. (b) River plot showing outgoing communication patterns of secreting cells for IDC.

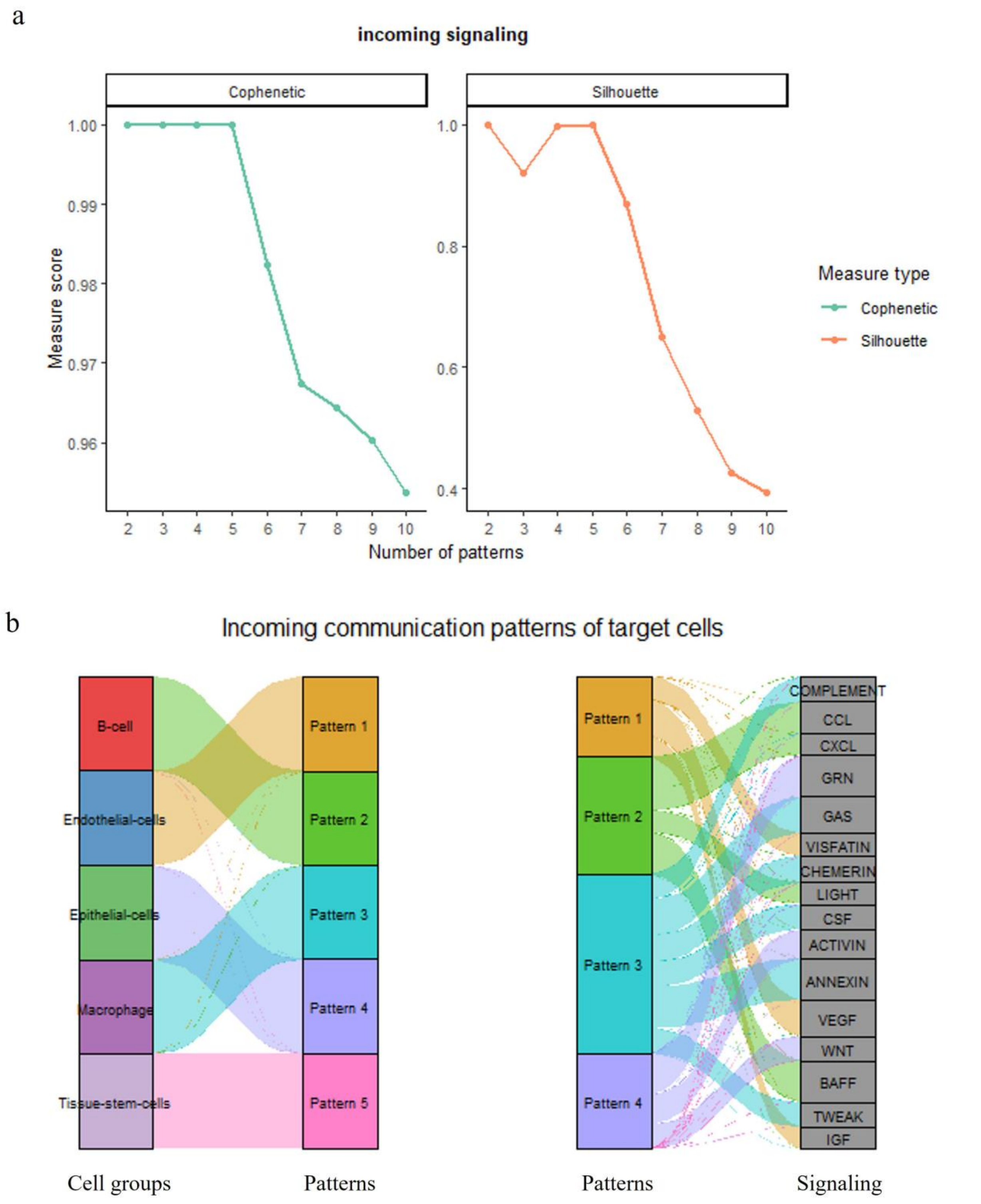


**Figure S14.** (a) Determination of the number of inferred incoming communication patterns for IDC. (b) River plot showing incoming communication patterns of secreting cells for IDC.

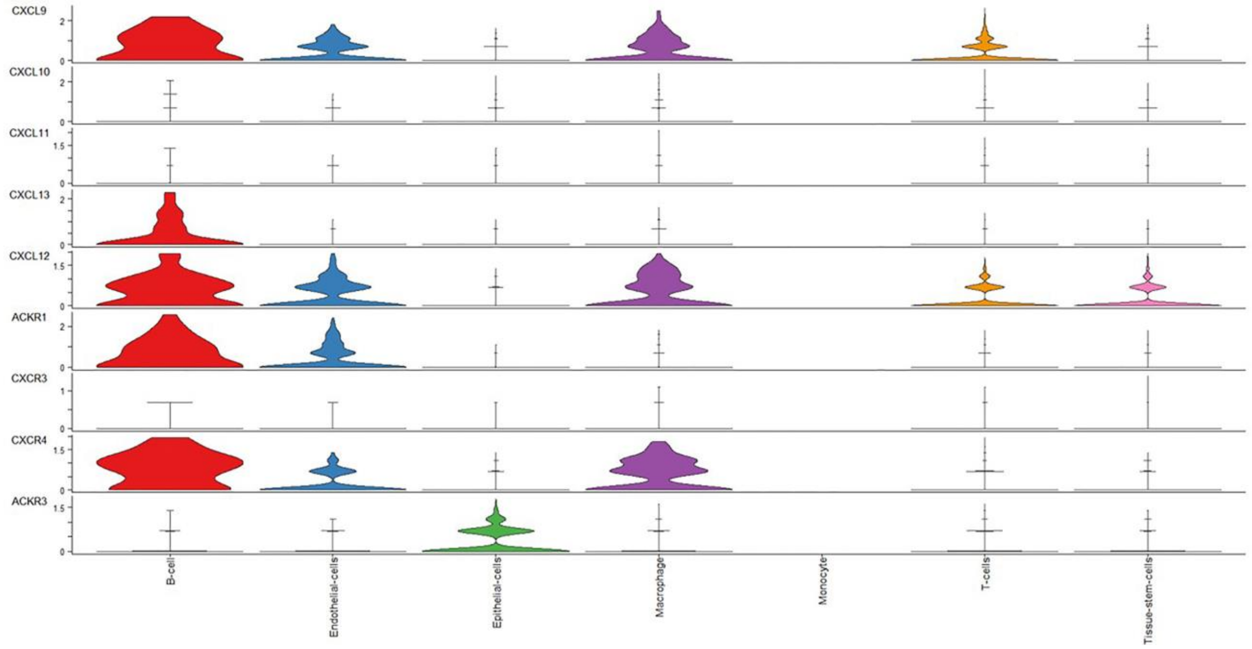


**Figure S15.** Gene Expression in CXCL Signaling Pathway for IDC.

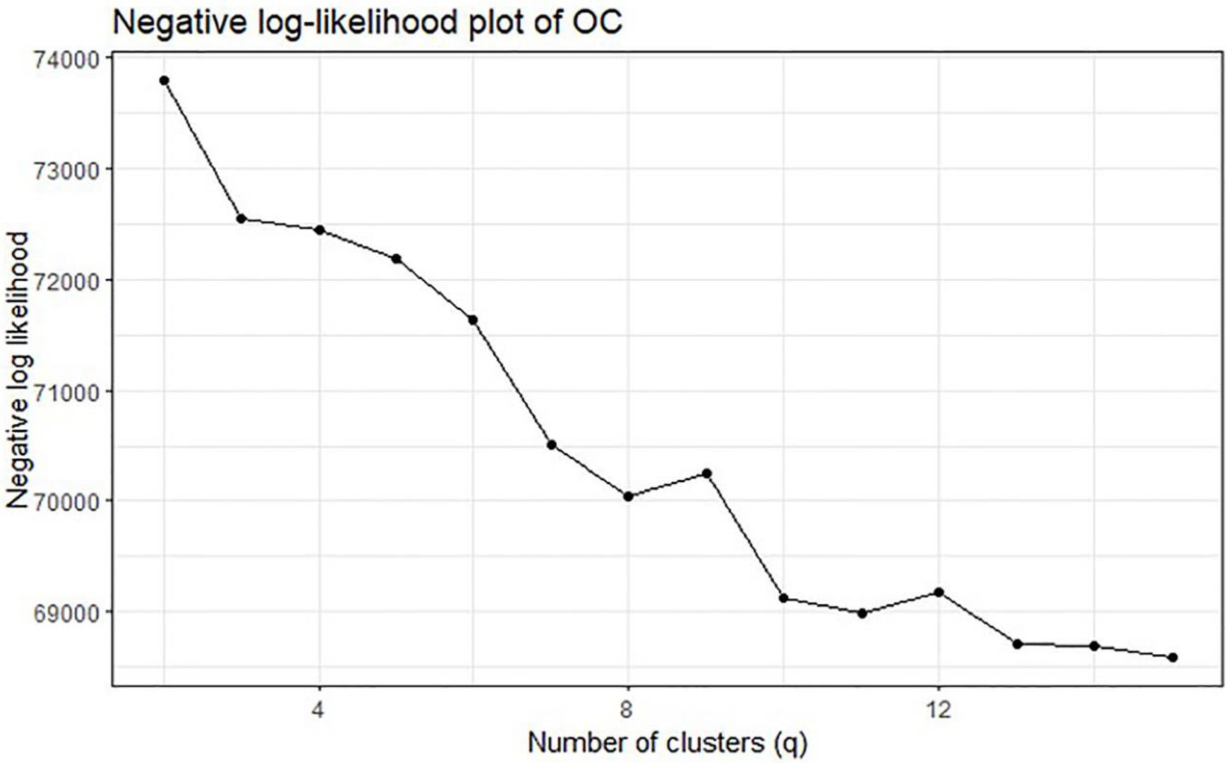


**Figure S16.** The optimal number of clusters for the OC dataset is determined to be 8 based on the elbow plot.

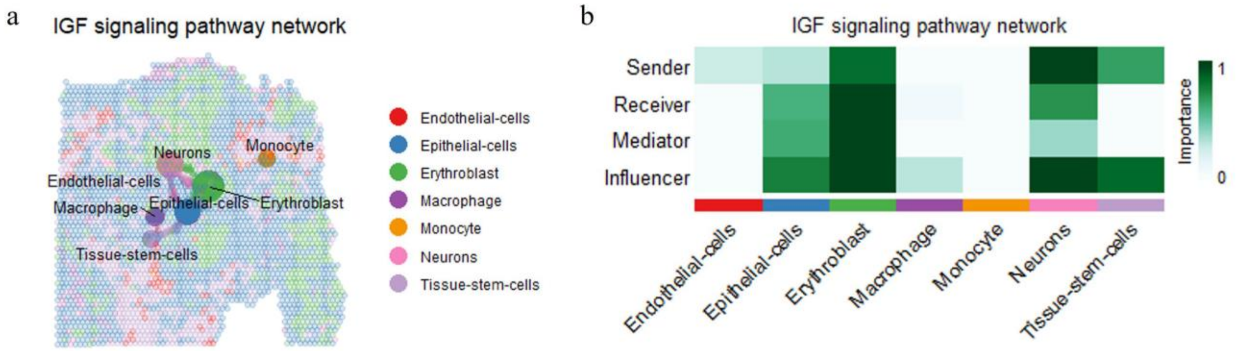


**Figure S17.** (a) Spatial map with signaling overlay of the IGF Signaling Pathway Network for OC. (b) Heatmap of Centrality Scores in IGF Signaling Pathway Network for OC.

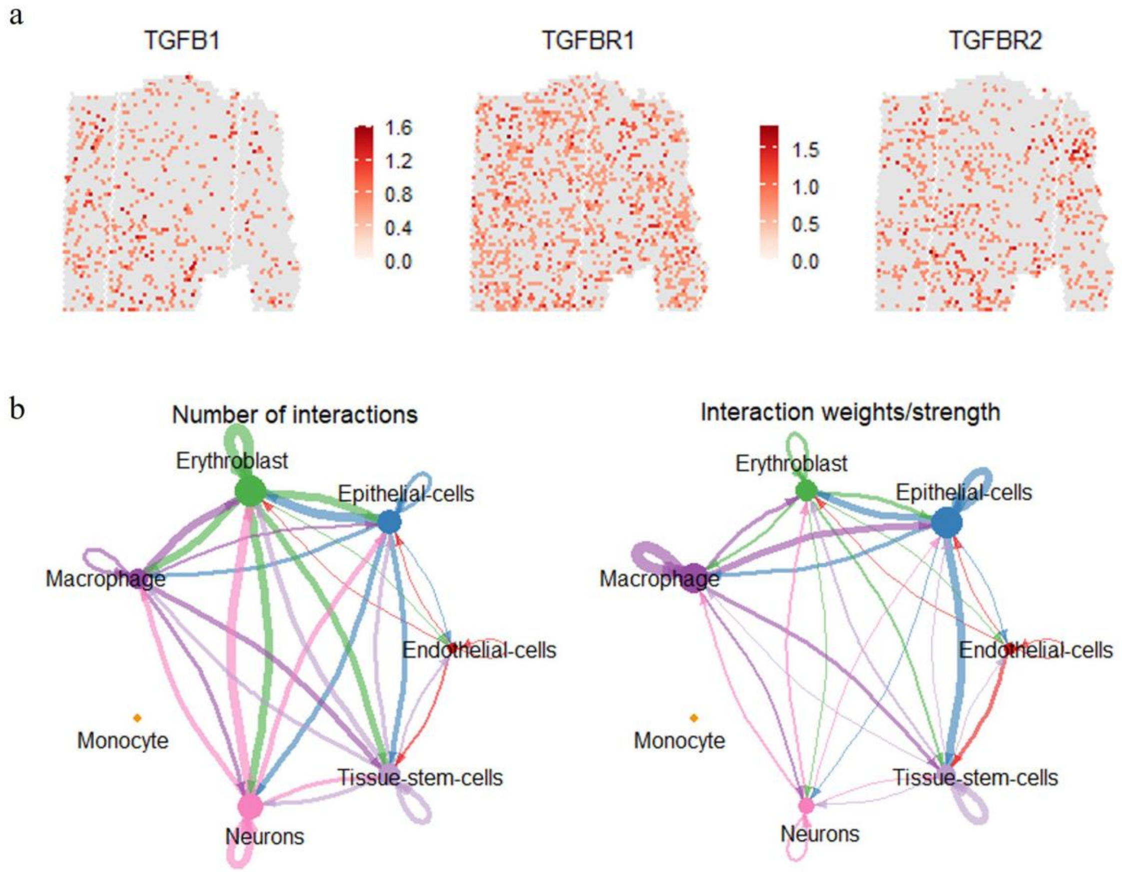


**Figure S18.** (a) Spatial Distribution of TGFB1, TGFBR1, and TGFBR2 Expression for OC. (b) Number of interactions and interaction weights for OC.

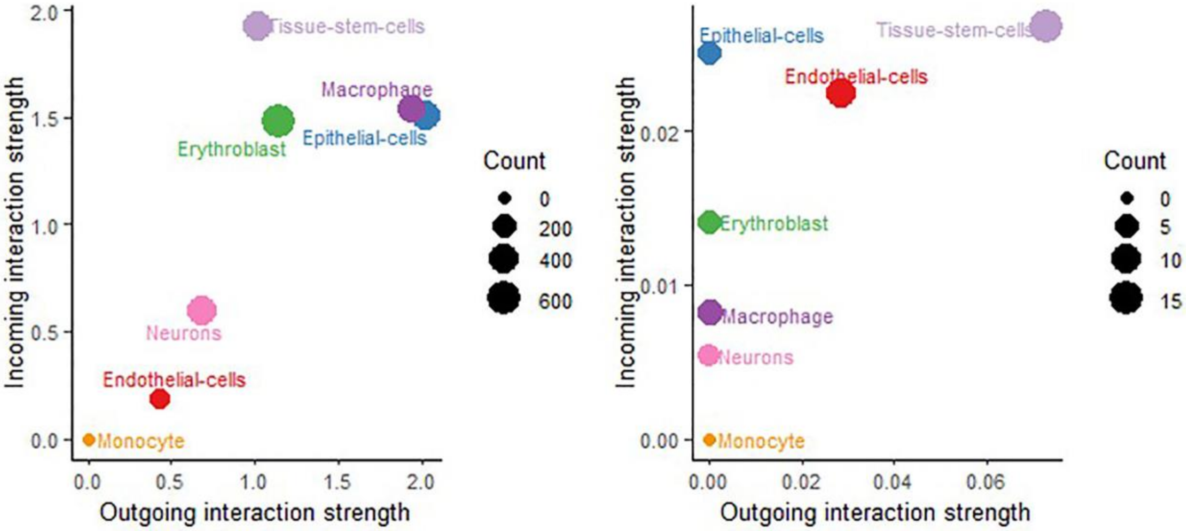


**Figure S19.** Global and CXCL/CCL- Specific Signaling Role Analysis in Cell-Cell Communication for OC.

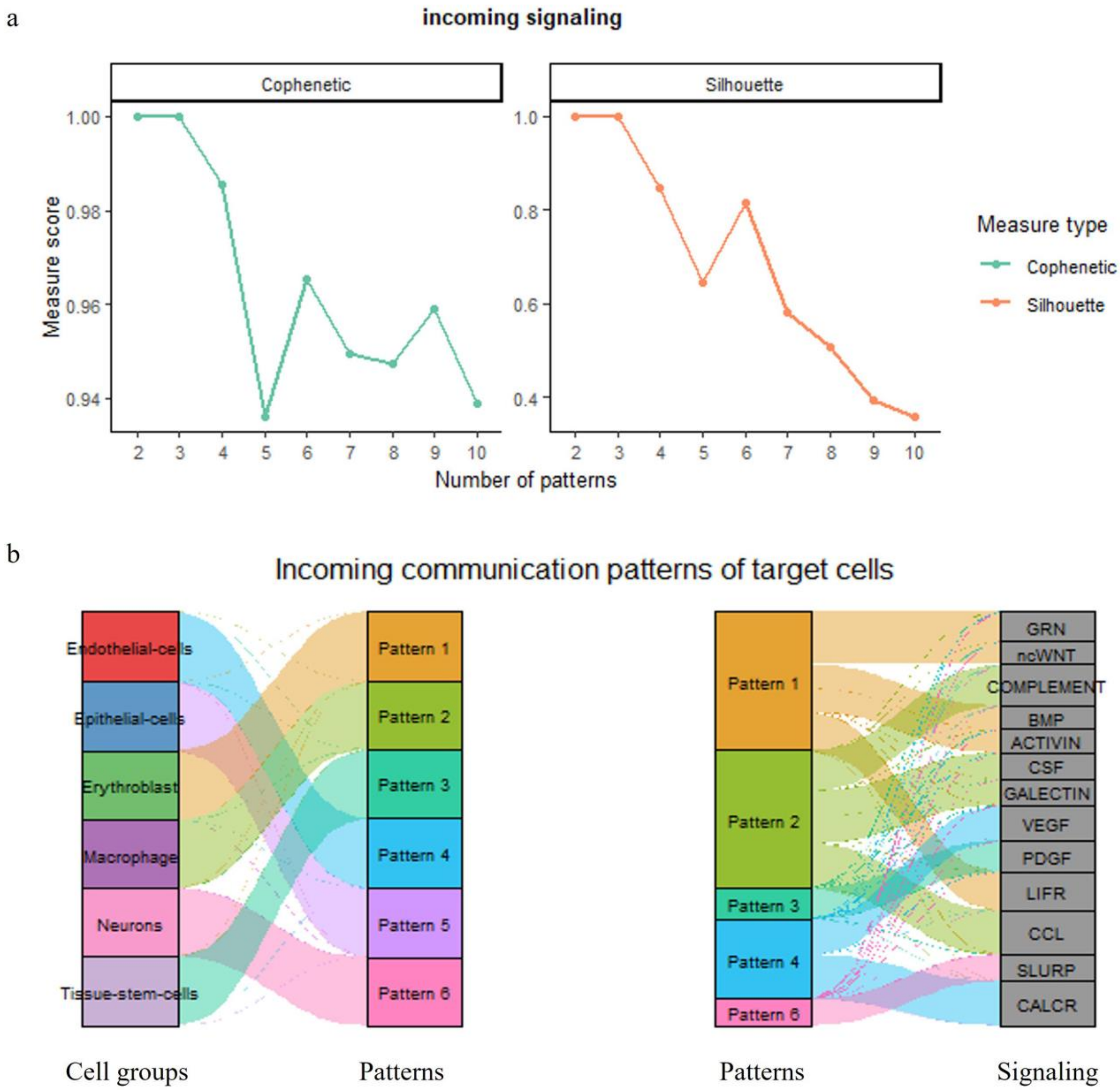


**Figure S20.** (a) Determination of the number of inferred incoming communication patterns for OC. (b) Incoming communication patterns of target cells for OC.

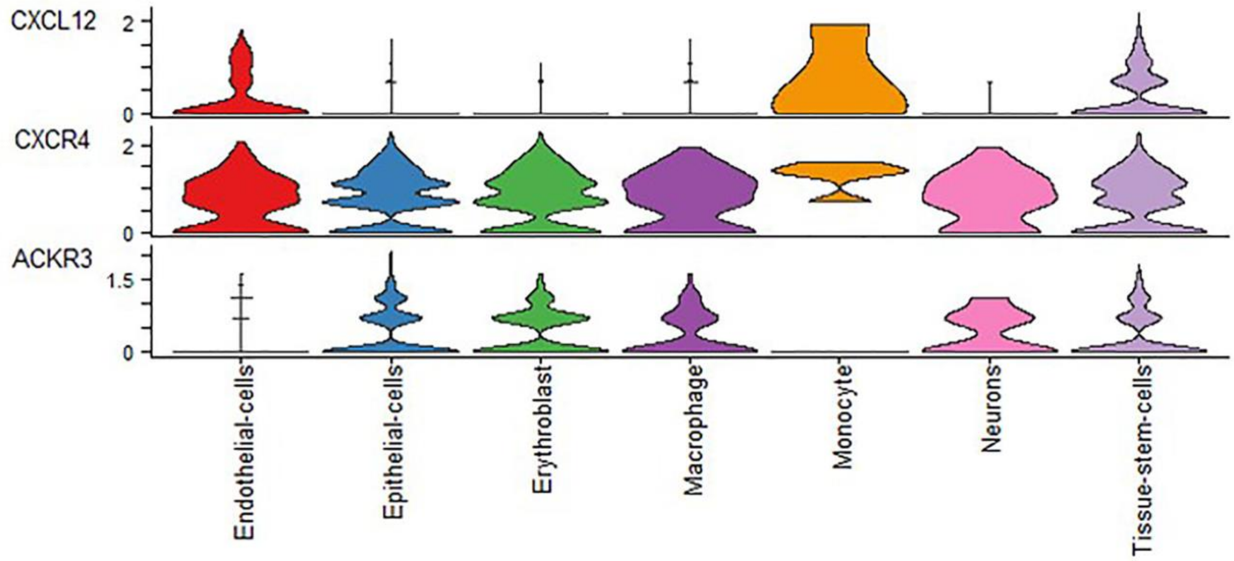


**Figure S21.** Gene Expression in CXCL Signaling Pathway for OC.

### 3. Supplementary Note:

#### 3.1 Supplementary Note 1: Application to the HER2 ST data

We applied SpatialESD to an eight-slice HER2 dataset which was manually annotated by Andersson [1]. The HER2 dataset was labeled into six categories: adipose tissue, cancer in situ, connective tissue, immune infiltrate, invasive cancer, and undetermined. In this analysis, we selected two base clustering methods, BASS and BayesSpace. The clustering performance of SpatialESD and the base methods was compared using the ARI, with detailed results presented in Table S5. The boxplot (Figure S22a) shows that SpatialESD achieved a slightly lower ARI value than BASS (by 0.016) but still outperformed other methods, exceeding them by 0.041 to 0.106. For spatial domain visualization analysis, we focused on slice C1, where the ARI value increased from the lowest base clustering result of 0.309 to 0.514 with SpatialESD, whereas the EnSDD ensemble approach resulted in a lower ARI of 0.297. Visualization results (Figure S22b-c) reveal that all base clustering methods performed poorly in identifying the invasive cancer region in the slice, particularly SpaGCN, which exhibited chaotic detection patterns. In contrast, SpatialESD consistently delivered superior visualization results compared to individual spatial domain detection methods as well as EnSDD.

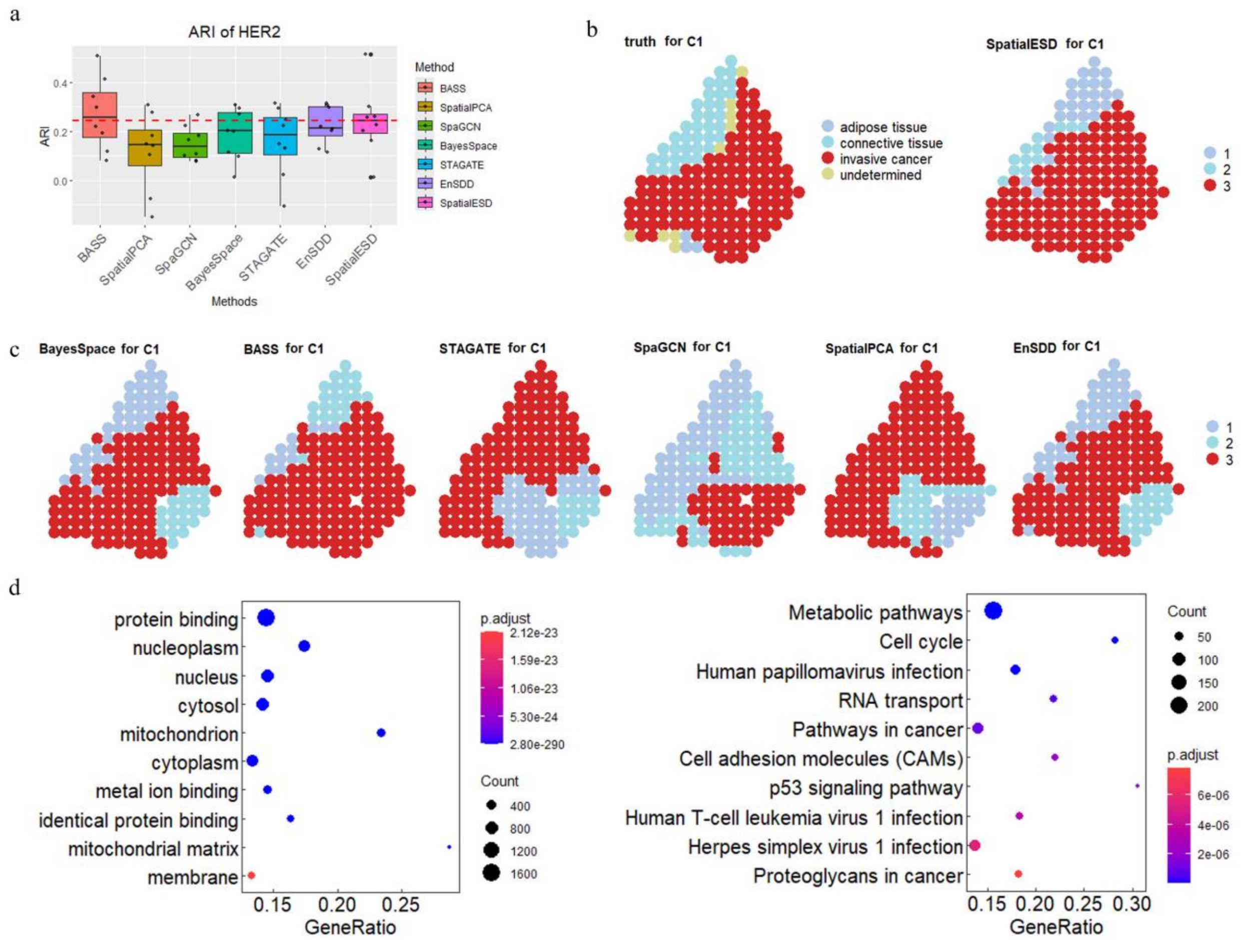


**Figure S22.** SpatialESD enhances tissue structure detection in HER2. (a) Boxplots of ARI. (b-c) Visualization of spatial domains using the SpatialESD method and several SDD methods. (d)

Enrichment analysis of differentially expressed genes (Left: GO enrichment; Right: KEGG pathway enrichment).

Based on the identified spatial domains by SpatialESD, the enrichment results (Figure S22d) revealed significant enrichment of spatially differentially expressed genes in multiple cellular components and molecular functions. The enriched cellular components include the nucleus, cytosol, mitochondrion, mitochondrial matrix, cytoplasm, membrane, and nucleoplasm. The molecular functions primarily include protein binding, metal ion binding, and identical protein binding. KEGG pathway enrichment analysis showed that the spatially differentially expressed genes were significantly enriched in several key pathways, including metabolic pathways, cell cycle, human papillomavirus infection, RNA transport, cancer-related pathways, cell adhesion molecules (CAMs), p53 signaling pathway, human T-cell leukemia virus 1 infection, herpes simplex virus 1 infection, and proteoglycans in cancer. The visual domain results for the eight tissue sections are shown in Supplementary Figure S11.

##### 3.2 Supplementary Note 2: Application to the Invasive Ductal Carcinoma ST data

We also performed corresponding trajectory analysis, where Figure S23a reveals the developmental direction from healthy tissue to tumor tissue, showing the developmental process of tumor cells and their interactions with immune cells in the tumor microenvironment.

Additionally, spatial cell-cell communication analysis was conducted (Figures S23b-e), where we integrated the ST data with the single-cell reference dataset HumanPrimaryCellAtlasData to explore the interactions between different cell types within the tissue (Figures S23c). The annotation results included seven cell types: B cells, endothelial cells, epithelial cells, macrophages, monocytes, T cells, and tissue stem cells. The analysis revealed significant spatial communication between tumor cells and immune cells, particularly in the tumor microenvironment, as well as interactions between tumor cells and stromal cells. Notably, we observed that Macrophage showed strong signaling interactions with B-cell (Figure S23b), which is consistent with previous research findings [2,3]. Figure S23d shows that there is a strong signaling connection between B cells, endothelial cells, and epithelial cells, reflected by the thicker connection lines between these cell types, indicating a strong interaction in their signaling pathways. In contrast, T cells, macrophages, and tissue stem cells show weaker connections with the above cell types, with thinner connection lines, suggesting less or weaker signaling interaction between these cells. Notably, there is no significant interaction between monocytes and other cell types, indicating that B cells, endothelial cells, and epithelial cells may play a more critical role in immune responses, cell migration, and other biological processes within the tissue, while their interactions with other cell types are relatively limited. Overall, this result suggests that B cells, endothelial cells, and epithelial cells may closely cooperate through a cellular communication network under specific physiological or pathological conditions, potentially playing an important role in immune responses and the tumor microenvironment. Figure S23d shows that the dark color of epithelial cells in all roles indicates their central role in the cell communication network, suggesting that they play an important role in signaling transmission, reception, mediation, and influence, particularly in processes such as immune response, cell migration, and tumor progression [4,5]. The first two plots in Figure S23e show the spatial distribution of TGFB1 (ligand)

and TGFBR1 (receptor) within the tissue, with a red gradient indicating the expression intensity of these molecules in different regions. The deeper the red, the stronger the expression. This result suggests the activity and spatial localization of TGFB1 and TGFBR1 within the tissue. The third plot further illustrates the expression of the TGFB1-TGFBR1-TGFBR2 ligand-receptor pair, specifically showing the signaling interaction between TGFB1 (ligand) and TGFBR1 and TGFBR2 (receptors). This plot allows us to observe the expression intensity and spatial distribution of these ligand-receptor pairs in the tissue, with deeper colors indicating stronger expression. It helps reveal the critical role of these molecules in cell-cell communication, especially their potential involvement in processes like immune response and tumor progression. This highlights the dynamic cell-cell interactions that shape the spatial architecture of the tumor and its surrounding microenvironment, providing valuable insights into the mechanisms of tumor progression and immune evasion. The complete results of the downstream analysis are provided in Supplementary Figures S12-S15.

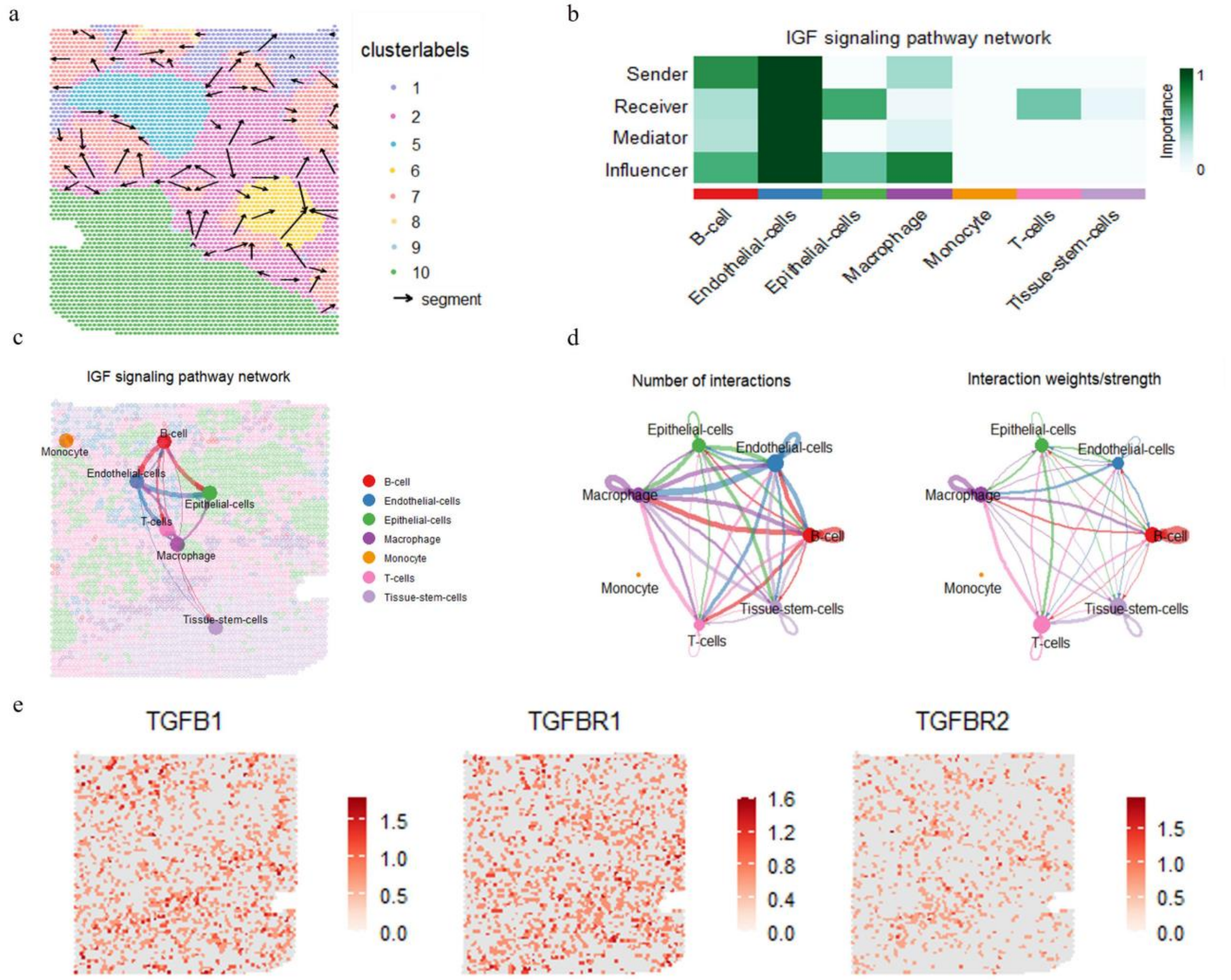


**Figure S23.** (a) Trajectory analysis for IDC. (b) Heatmap of Centrality Scores in IGF Signaling Pathway Network for IDC. (c) Spatial map with signaling overlay of the IGF Signaling Pathway Network for IDC. (d) Number of interactions and interaction weights for IDC. (e) Spatial Distribution of TGFB1, TGFBR1, and TGFBR2 Expression for IDC.

##### 3.3 Supplementary Note 3: Application to the Ovarian Cancer ST data

Trajectory analysis (Figure S24a) indicated migration or infiltration of CD45-labeled immune cells into other regions, suggesting their dynamic role in shaping tumor progression and immune responses within the TME.

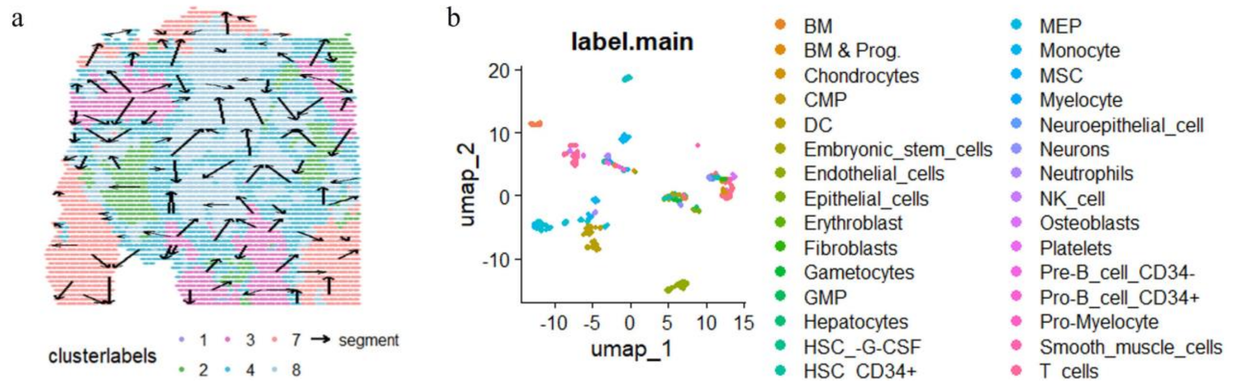


**Figure S24.** (a) Trajectory analysis for OC. (b) Integration of the OC dataset with single-cell data.

Additionally, we performed spatial cell-cell communication analysis (Figures S25 and Figures S26), integrating spatial transcriptomic data with a single-cell reference dataset (Figure S24b) to explore interactions among distinct cell types within the tissue. The annotated cell types included seven categories: erythroblasts, endothelial cells, epithelial cells, macrophages, monocytes, neurons, and tissue stem cells. The analysis of communication quantity (Figure S25, upper) and interaction strength (Figure S25, lower) revealed that erythroblasts exhibited the highest number of interactions, suggesting their frequent cell-cell communication within the local microenvironment, while macrophages displayed the strongest interaction strength, implying stable or intense signal exchange potentially related to inflammatory signaling, phagocytic regulation, or immune microenvironment shaping. Erythroblasts exhibited the highest number of cell-cell interactions, highlighting their active communication within the TME. Macrophages displayed robust signal exchange, underscoring their pivotal role in tumor immunity, particularly tumor-associated macrophages (TAMs) in driving tumor progression and immune evasion [6].

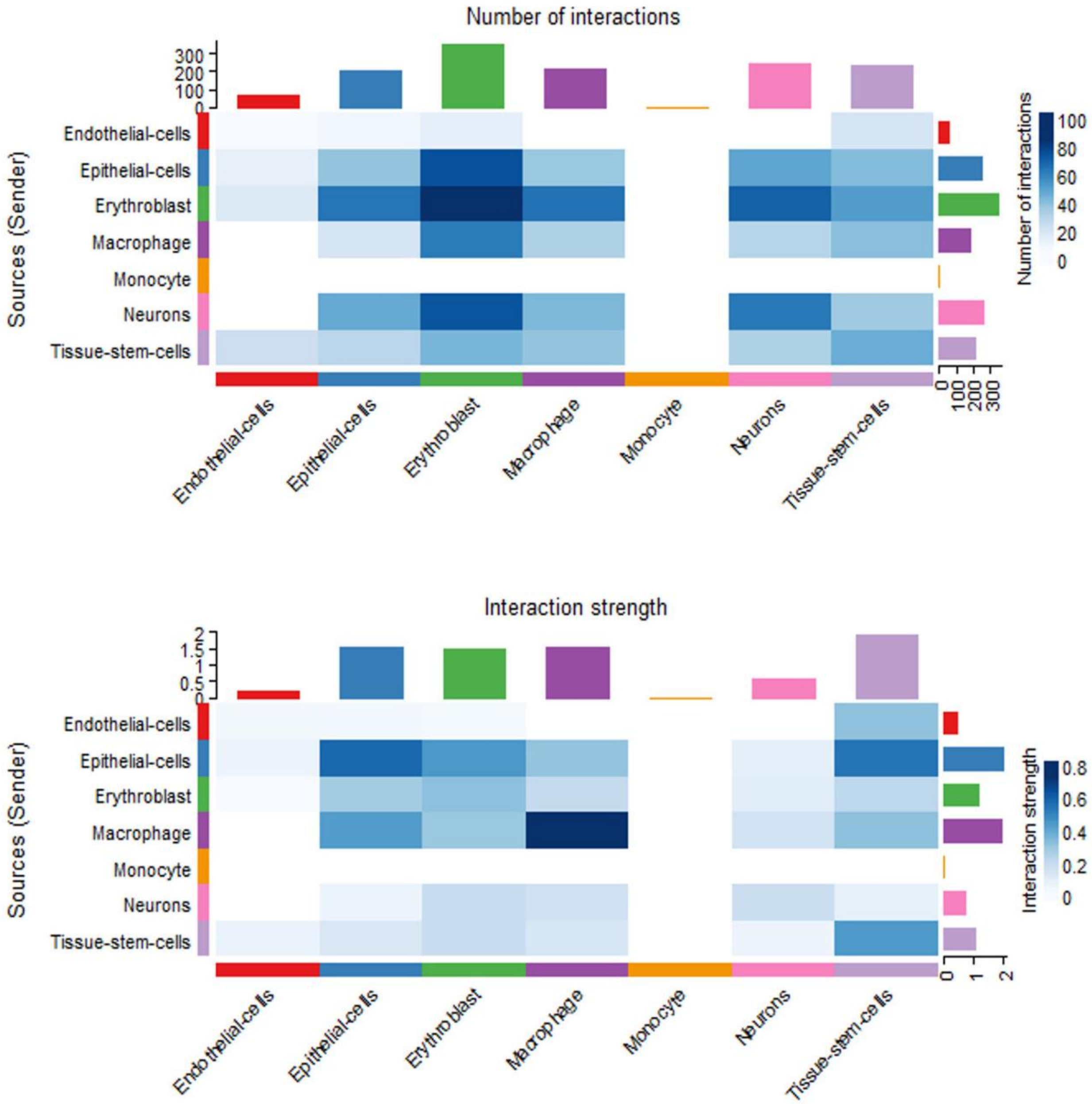


**Figure S25.** Number of Interactions and Interaction Strength for OC.

For outgoing communication patterns (Figure S26a), both the Cophenetic and Silhouette values showed abrupt declines when the number of output patterns reached 6. Cell pattern analysis (Figure S26b, left) demonstrated high similarity in communication patterns across different cell types, with closely branched structures, indicating shared signaling characteristics. Contribution analysis (Figure S26b, right) revealed distinct cell-type dominance in specific patterns: endothelial cells were most active in Pattern 6, epithelial cells enriched in Pattern 4, macrophages concentrated in Pattern 2, and erythroblasts dominated Pattern 1. Communication pattern analysis highlighted pathway-specific contributions: the CALCR pathway dominated Pattern 4 (darkest color), ANGPTL, ILT, and SLURP pathways contributed most to Pattern 2, while PTN, PARs, and VEG pathways played central roles in Pattern 1. Cellular pattern analysis revealed shared signaling features among distinct cell types, with endothelial cells predominantly contributing to Pattern 6, epithelial cells to Pattern 4, macrophages to Pattern 2, and erythroblasts to Pattern 1. Pathway-specific dominance was observed across patterns: CALCR signaling dominated Pattern 4 (epithelial cells), ANGPTL, ILT, and SLURP pathways in Pattern 2 (macrophages), and PTN, PARs, and VEG pathways in Pattern 1 (erythroblasts), implicating these pathways in angiogenesis and immune regulation. These findings provide novel insights into the cellular communication mechanisms within the tumor immune microenvironment, offering a theoretical foundation for future immunotherapeutic strategies. A River plot (Figure S26b) visually summarized the distribution of cell types across communication patterns and the pathway contributions within each pattern. The complete results of the downstream analysis are provided in Supplementary Figures S16-S21.

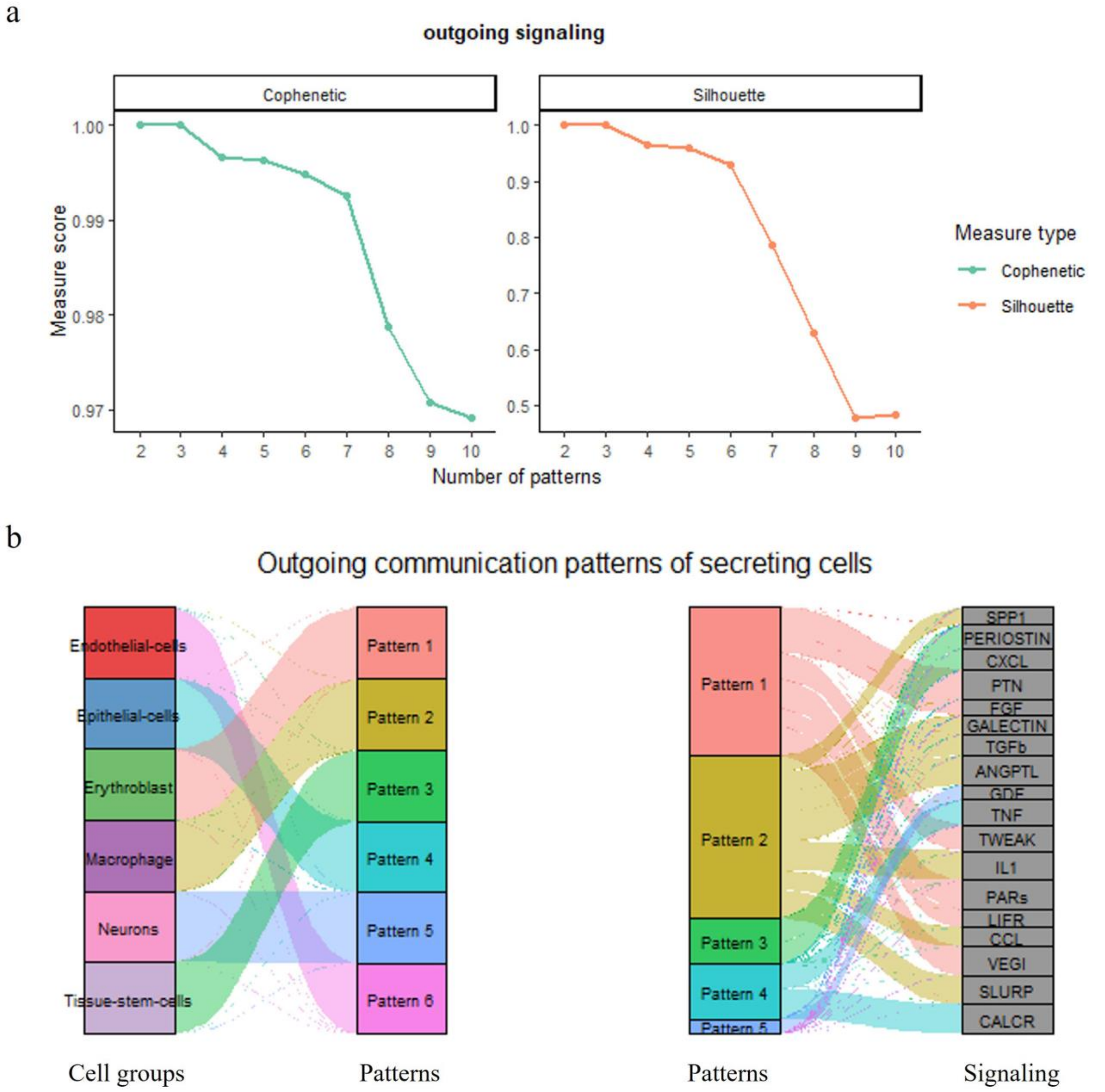


**Figure S26.** (a) Determination of the number of inferred outgoing communication patterns for OC. (b) River plot showing outgoing communication patterns of secreting cells for OC.
